# Human Regnases are antiviral restriction factors targeting viral RNA

**DOI:** 10.64898/2026.07.06.736706

**Authors:** Linda Grabe, Sarah Hommel, Lucas Singer, Maddalena Zangari, Kerstin Regensburger, Asimenia Vlachou, Rayhane Nchioua, Konstantin Sparrer, Mateusz Wawro, Aneta Kasza, Daniel Sauter, Frank Kirchhoff, Dorota Kmiec

**Affiliations:** Institute of Molecular Virology Ulm University Medical Center Ulm, Germany; German Center for Neurodegenerative Diseases (DZNE) Ulm, Germany; University Medical Center Hamburg-Eppendorf (UKE) Hamburg, Germany; Department of Cell Biochemistry Faculty of Biotechnology, Biochemistry and Biophysics Jagiellonian University Krakow, Poland; Institute for Medical Virology and Epidemiology of Viral Diseases University Hospital Tübingen Tübingen, Germany

## Abstract

Regnases regulate immune gene expression by degrading cellular mRNAs. Regnase-1 also targets viral RNA, but whether this activity is shared among other human Regnases remains unknown. Here, we systematically compared the antiviral properties of all four Regnases. Expression of Regnases-1–4 inhibited RNA viruses HIV-1, HIV-2, MLV, RSV, OC43 and SARS-CoV-2, but not DNA virus HSV-1. Depletion experiments demonstrated that endogenous Regnases restrict HIV-1 replication in a cell-type-dependent manner, with Regnase-1 and Regnase-4 exerting the strongest effects in T-cells and monocytes, and Regnase-2 and Regnase-3 in primary macrophages. All Regnases displayed signatures of positive selection, consistent with their roles as antiviral restriction factors, but only Regnase-1 and -4 were induced by interferons. Mechanistically, the antiviral activity of Regnases required an intact catalytic core, a CCCH zinc finger, and, in the case of Regnase-1, -2 and -3, a dimerisation motif (P/R). Chimeric Regnase-1 and Regnase-3 constructs suggested that differences in antiviral potency primarly reflect differences in RNA recognition rather than catalytic activity. Bicistronic reporter assays suggested that Regnase-1 and -4 exhibit broad targeting, whereas Regnases-2 and -3 display greater substrate selectivity and correlate in target specificity. Collectively, our findings establish Regnases as antiviral restriction factors with distinct viral RNA targeting specificities.

## INTRODUCTION

Intrinsic immunity is composed of a set of cell-autonomous defences that act directly on the virus and thereby determine the outcome of the infection, serving as an important first line of defence against pathogens ^1^. So-called antiviral restriction factors are a key component of this cell-intrinsic immunity and comprise a group of structurally diverse cellular proteins that inhibit viruses at different steps of their replication cycle. Viral RNA and transcription are well-established targets of many restriction factors such as APOBEC3 proteins, OAS proteins, SAMHD1, IFI16 and Zinc-Finger Antiviral Protein (ZAP) ^1–3^. It has become increasingly clear that multiple host ribonucleases such as RNase L, N4BP1 and KHNYN also participate in antiviral defence by targeting viral RNAs for decay ^4–10^. The latter two proteins are related to the human Regnase family, which has recently attracted considerable interest due to its broad immunomodulatory roles in innate immunity ^11–13^. MCPIP1-4, also known as ZC3H12A-D or Regnases 1-4, are PIN-domain ribonucleases with established roles in post-transcriptional control of inflammatory gene expression and immune homeostasis. Their well-described immune functions and pre-existing evidence that Regnase-1 inhibits multiple RNA viruses ^14–19^ suggest that, in addition to their immunomodulatory roles, Regnases might act as antiviral restriction factors. While Regnase-1 is well established as a direct antiviral effector ^14–22^, evidence supporting the antiviral roles for Regnases-2–4 is lacking. In resting CD4+ T cells, Regnase-1/ZC3H12A/MCPIP1 restricts HIV-1 infection and is rapidly degraded by MALT-1 upon T-cell activation^17^. These findings support the view that Regnase-1 represents a barrier to HIV-1 infection, which the virus overcomes by preferentially replicating in activated CD4+ T cells. Regnase-1 has also been reported to inhibit viruses such as hepatitis B (HBV) and C (HCV) and coxsackievirus B3 through binding and cleavage of viral RNA ^15,16,18^. In the case of HBV, the viral epsilon RNA was found to be a determinant of susceptibility, suggesting that Regnase-1 specifically recognises structured viral RNA ^14^. However, much of the literature on the antiviral functions of Regnase-1 relies on overexpression systems, while evidence of its endogenous function is limited. Also, Regnase-1 does not act only on viral substrates. Rather, it acts as a multifunctional RNase that targets viral RNAs while also reshaping the host transcriptome by degrading multiple cellular transcripts predominantly involved in inflammation and immune response homeostasis ^11,13,23,24^.

While Regnase-1 has emerged as an antiviral factor, the interplay between other members of the Regnase family and viruses is poorly understood. Regnases-2, -3, and -4 share the catalytic PIN/NYN domain and CCCH zinc finger responsible for Regnase-1’s activity, and could therefore, in principle, target viral RNA. However, they have been studied almost exclusively as regulators of inflammatory gene expression ^25–28^. The experiments needed to establish physiological restriction, such as binding and cleavage of viral RNA or loss of viral control upon depletion of the endogenous protein, have not been reported for Regnases-2 to -4. Regnase-3 has been linked to the regulation of interferon homeostasis in myeloid cells ^11,24,29,30^. Thus, it remains to be established if the Regnases represent a functionally conserved class of antiviral restriction factors that target viral RNA.

To understand how RNA decay by Regnase family proteins intersects with the antiviral response, we comprehensively compared the antiviral potential of human regnases at both overexpression and endogenous levels, and uncovered the molecular determinants of their activity. The analysis uncovered a broad antiviral mechanism against RNA viruses, with differences in expression and target specificity that shape the effectiveness of human Regnases as antiviral restriction factors in different cell types.

## MATERIALS AND METHODS

### Ethical statement

Experiments involving cells derived from human blood were reviewed and approved by the Institutional Review Board (i.e. the Ethics Committee of Ulm University, vote number 50/16). Individuals and/or their legal guardians provided written informed consent prior to donating blood. All human-derived samples were anonymized before use. The use of established cell lines did not require the approval of the Institutional Review Board.

### Virus strains

HSV-1-GFP (F-strain) was kindly provided by Prof. Benedikt Kaufer, FU Berlin, Berlin, Germany. RSV-GFP was kindly provided by Dr. Kariem Ezzat Ahmed, Karolinska Institute, Stockholm, Sweden; and Dr. Michael N. Teng, University of South Florida, Tampa, USA^31^. SARS-CoV-2 BetaCoV/Netherlands/01/NL/2020 was obtained from the European Virus Archive and HCoV-OC43 was obtained from American Type Culture Collection (ATCC; CR-1558TM). Infectious molecular clones of HIV and SIV were described before ^32–37^.

### Cell lines and culture

HEK293T, HEp-2 and HeLa cells were obtained from the American Type Culture Collection (ATCC). Generation of CEM-M7 Cas9 has been described before ^37^. THP-1-Cas9 cells were purchased from Applied Biological Materials Inc. U373 MAGI cells were obtained from the NIH AIDS Reagent Program and were transduced with a Cas9-expressing lentiviral vector as described before ^37,38^. TZM-bl cells, which express CD4, CCR5, and CXCR4 and contain the β-galactosidase genes under the control of the HIV promoter, were obtained from the NIH AIDS Reagent Program ^39,40^.

HEK293T, HEp2, U373 MAGI, HeLa and TZM-bl reporter cells were cultured in Dulbecco’s modified Eagle medium supplemented with 10% fetal calf serum (FCS), 100 U/ml penicillin and 100 μg/ml streptomycin. CEM-M7-Cas9 and THP-1-Cas9 were cultured in Roswell Park Memorial Institute (RPMI) 1640 Medium supplemented with 10% FCS, 100 U/ml penicillin and 100 μg/ml streptomycin, with an additional 1.2 μg/ml puromycin for THP-1-Cas9 cells.

All cells were grown at 37°C in a humidified atmosphere with 5% CO_2_, except for OC43-infected cultures, which were incubated at 32°C.

### Isolation and stimulation of primary cells

PBMCs were isolated from human buffy coats (DRK, Ulm) using human Pancoll (PAN-Biotech) and gradient density centrifugation as described before (https://www.protocols.io/view/peripheral-blood-mononuclear-cell-isolation-and-st36wgq9rkolk5/v4).

To generate MDMs, 4 × 10^6^ PBMCs/ mL were seeded in RPMI supplemented with 100 µg/mL streptomycin, 100 U/mL penicillin, 10% human AB serum (PeproTech), and 15 ng/mL M-CSF (PeproTech). Cells were cultured until differentiation (6-9 days), with medium changes every 3 days. Non-adherent cells were removed by washing with PBS.

CD4^+^ T cells were isolated from human buffy coats (DRK, Ulm) using the RosetteSep Human T Cell Enrichment Cocktail (Stemcell Technologies) according to the manufacturer’s recommendations. For experiments, 4 × 10^6^ isolated CD4^+^ T cells/mL were cultured in 12-well plates in RPMI medium supplemented with 10% FCS, L-glutamine (2 mM), streptomycin (100 μg/mL) and penicillin (100 U/mL).

To check mRNA induction, cells were treated with either IFNα (500 U/mL), IFNβ (500 U/mL), and IFNγ (200 U/mL), or left untreated and harvested after 2-72 h (as specified in figure legends).

### siRNA knockdown

Fully differentiated human MDMs were transfected with siRNA twice, with a resting day in between. Cells were transfected using Regnase-specific (Dharmacon #L-014576-01-0010; L-024828-02-0010; L-026513-01-0010; L-024742-01-0010) or non-targeting pool siRNA (Dharmacon #D-001810-10-20) using Lipofectamine RNAiMax (Life Technologies) according to the manufacturer’s instructions. One day after the second knockdown, the cells were infected with HIV-1 AD8 virus stocks overnight. The following day, the input virus was removed, cells were washed in PBS, and fresh medium was added. Virus-containing supernatants and cells were harvested at 3-4 dpi. Infectious virus yield was determined using the TZM-bl infectivity reporter assay, and the knockdown efficiency was determined using qRT-PCR.

### Expression constructs

The pcDNA4-3xFLAG expression vectors encoding wild-type Regnase-1–4 were generated using restriction cloning from previously described templates^25,28,41,42^. The resulting PCR products were digested with NotI/XbaI (for Regnase-1) or NotI/XhoI (for Regnase-2–4) and ligated into the pcDNA4-3xFLAG backbone digested with the corresponding restriction enzymes. The pCG N-FLAG IRES BFP Regnase1-4 expression vectors were generated by subcloning the Regnase inserts from the pcDNA vector into pCG using Gibson assembly (NEB) and XbaI/MluI restriction sites. Mutants affecting PIN domain catalytic activity, dimerisation and RNA binding by ZnF were generated to assess their contribution to antiviral activity. Dimerisation-deficient mutants were generated by introducing amino acid substitutions corresponding to the previously described Regnase-1 mutations P212A and R214A, which are predicted to disrupt PIN domain dimerisation ^43^. The corresponding positions in Regnase-2-4 were identified by sequence alignment and mutated accordingly. For ZnF mutants, the motif was targeted by substituting CCCH residues with alanines (A). Primers were synthesized by Biomers.net. Site-directed mutagenesis was performed using the Q5 site-directed mutagenesis Kit (NEB). All constructs were verified by Sanger Sequencing.

**Table 1.**
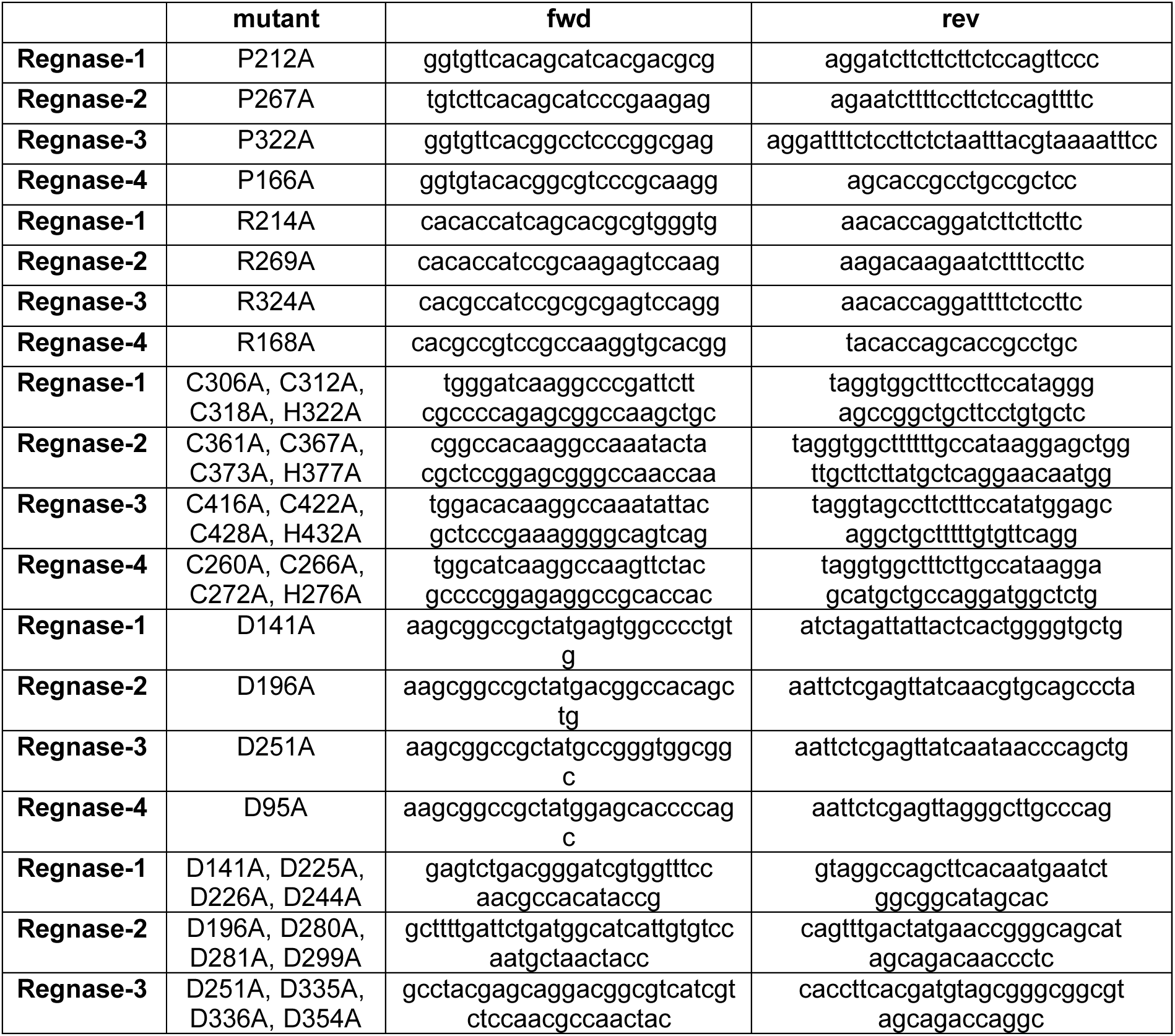
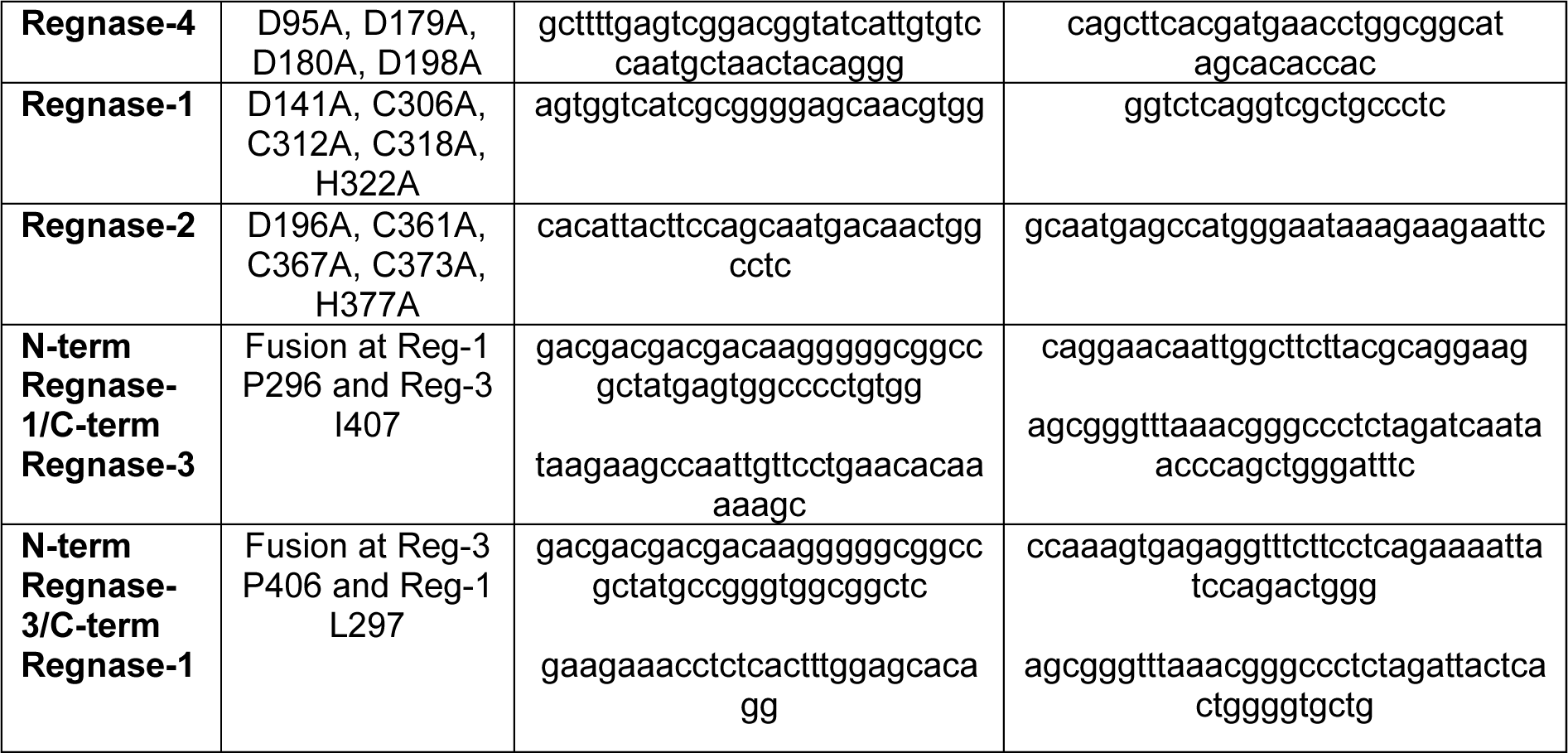
Primer sequences used for Regnase mutagenesis.

pmirGLO-3’UTR mouse IL-6 was a gift from Silvia Monticelli (Addgene plasmid # 207126) and has been described before ^44^. HIV-1 NL4-3 LTR luciferase reporter construct was described before ^45^. Mutations within the TAR loop were generated using the Q5 site-directed mutagenesis kit (NEB) and the primers: TAR deletion (ccagtacaggcaaaaagcagctgcttatatgcagcatc;cactgcttaagcctcaataaagcttgccttgagtg), scrambled TAR (ctgtacacttttgactgcgcgtcaccagtacaggcaaaaagcagctgc; ccttctcgtgagtggttcgtgccagcatcctgcactgcttaagcctcaataaagcttg). To generate the control plasmid, the HIV-1 LTR was replaced with a CMV promoter using Gibson assembly (NEB) and the following primers (ggtaccgagctcttacgcgtgacattgattattgactagt;gtaccggaatgccaagcttacttagatcgcagatctcgaagctctgcttatataga cc).

### Generation of individual sgRNA HIV-1 NL4-3 constructs

The HIV-1 NL4-3 proviral backbone containing an sgRNA expression cassette comprising the human U6 promoter has been described before^37^. To generate individual CRISPR-competent HIV-1 NL4-3 constructs, sgRNAs targeting Regnase1–4, zinc finger antiviral protein (ZAP) as positive control, as well as a non-targeting (NT) control, were cloned by Gibson cloning using BsmBI vector digestion and complementary oligonucleotides synthesized by Biomers.net (Reg-1 sgRNA 1: tcttgtggaaaggacgaaacaccggagtggaagcgcttcatcggttttagagctagaaatagcaag; Reg-1 sgRNA 2: tcttgtggaaaggacgaaacaccgaaggaggtcttctcctgccggttttagagctagaaatagcaag; Reg-2 sgRNA 1: tcttgtggaaaggacgaaacaccgatgatccattaggacgccagttttagagctagaaatagcaag; Reg-2 sgRNA 2: tcttgtggaaaggacgaaacaccgaacctcagaggtcaaacgtggttttagagctagaaatagcaag; Reg-3 sgRNA 1: tcttgtggaaaggacgaaacaccgtatggataccgttaatgtggttttagagctagaaatagcaag; Reg-3 sgRNA 2: tcttgtggaaaggacgaaacaccgtcaaacgaggtgctccaaaggttttagagctagaaatagcaag; Reg-4 sgRNA 1: tcttgtggaaaggacgaaacaccgtgctcagggagggtccatgggttttagagctagaaatagcaag; Reg-4 sgRNA 2: tcttgtggaaaggacgaaacaccgcacccaggctagtgcctcggttttagagctagaaatagcaag; ZAP sgRNA 1: tcttgtggaaaggacgaaacaccgtctggtagaagttatatctggttttagagctagaaatagcaag; NT: tcttgtggaaaggacgaaacaccgacggaggctaagcgtcgcaagttttagagctagaaatagcaag). sgRNA inserts were assembled into the linearized backbone using NEBuilder® HiFi DNA Assembly Master Mix (NEB) according to the manufacturer’s instructions and transformed into chemically competent *E. coli* Q5 strain (NEB). Plasmid DNA was purified using the Wizard® Plus Midipreps DNA Purification System (Promega) following the manufacturer’s protocol. Correct sgRNA insertion was confirmed by Sanger sequencing (Eurofins).

### Assessment of CRISPR/Cas9 knockout efficiency

For CRISPR/Cas9-mediated gene knockout, 1 × 10⁶ CEM-M7-Cas9 cells were electroporated with HiFi Cas9 Nuclease V3 (IDT) pre-complexed with sgRNAs (80 pmol Cas9 and 300 pmol sgRNA; Lonza). Cells were transfected using the Amaxa 4D-Nucleofector™ system (P3 Primary Cell 4D-Nucleofector™ X Kit S, Cat.#V4XP-3032, Lonza) with pulse code EO-115 with the following sgRNAs:

Regnase-1 (sgRNA 1: ggagtggaagcgcttcatcg; sgRNA 2: aaggaggtcttctcctgccg); Regnase-2 (1: gatgatccattaggacgcca; 2: aacctcagaggtcaaacgtg); Regnase-3 (1: gtatggataccgttaatgtg; 2: tcaaacgaggtgctccaaag); Regnase-4 1: tgctcagggagggtccatgg; 2: gcacccaggctagtgcctcg); Non-targeting control (acggaggctaagcgtcgcaa).

At four days post-transfection, cells were harvested for knockout analysis. Genomic DNA was isolated using the QIAamp® DNA Mini Kit (Qiagen). Editing efficiency at the genomic level was determined by PCR amplification of a ∼300 bp region spanning the CRISPR cut site using PCR and the following primers: Reg-1 sgRNA 1 (cctgcgggaactggagaagaag; gcaacctcccatggccacag), Reg-1 sgRNA 2 (ctgctcactaccaccaccaaag; gctggggcagtaagaccatcc), Reg-2 sgRNA 1 (gcagaggatagacttgaggcag; ctcaaagacagtggacaagcc), Reg-2 sgRNA 2 (caagtgcaaatactaccatcc; cacttagggcagtcacaagc), Reg-3 sgRNA 1 (ggagtcaagtacacgtaacaac; cctgcagagctgtctgtgag), Reg-3 sgRNA 2 (ggtcagtggctgatgaactcc; gcagtagaagtagctgggatg), Reg-4 sgRNA 1 (cgaatgcttcctggtgttcttg; gagggcagaacttgggcaagag), Reg-4 sgRNA 2 (cttccagaagctgggctatg; gctacaggatgaagggaagc), ZAP sgRNA 1 (ctgcaaacacctgaagctgtg; ctcttgggtcagcatcatctg), ZAP sgRNA 2(ctggtatagatggaacctgtc; cttccctttgataatgttgg), followed by Sanger sequencing. Sequencing results from KO samples were compared to the corresponding non-targeting (NT) control sequence using the Tracking of Indels by DEcomposition (TIDE) algorithm (https://tide.nki.nl)^46^, which aligns the sgRNA target region to the control sequence to define the expected Cas9 cleavage site and decomposes the sequencing trace downstream of the cut site into a mixture of indel-containing sequences of varying lengths. Total knockout efficiency was defined as the overall indel percentage reported by the TIDE algorithm, corresponding to the sum of all detected indel frequencies within the decomposition window.

### Transfection of cells

HEK293T cells (0.4 × 10⁶ cells/mL) were seeded and transfected using PEI (3:1 PEI to DNA ratio) with 0.5 µg HIV-1 proviral DNA and between 0-0.5 µg pcDNA4 or pCG protein expression construct. The total amount of DNA was normalised to 1 µg using pCG or pcDNA-based GFP expression vector. Media were replaced 24 h post-transfection, and cell-free virus-containing supernatants and transfected cells were harvested 48 h post-transfection. To determine infectious virus production, 1 × 10^4^ TZM-bl cells per well were seeded in 96-well cell culture plates and infected in triplicate with the virus-containing supernatants. Viral infectivity was assessed 48-72 h post-infection using the Gal-Screen® kit (Applied Biosystems) and Orion microplate luminometer (Berthold).

### Production of HIV-1 virus stocks

HEK293T cells were seeded in 6-well plates at a density of 0.4 × 10⁶ cells/mL and cultured overnight. The following day, cells were transfected with 5 µg of HIV-1 AD8 (for MDMs) or HIV-1 NL4-3 sgRNA constructs with or without VSV-G expression plasmid using TransIT-LT1 Transfection Reagent (Mirus) at a ratio of 3 µL transfection reagent per 1 µg plasmid DNA, according to the manufacturer’s instructions. At 24 h post-transfection, the culture medium was replaced. Cell-free virus-containing supernatants were harvested 24 h later and clarified by centrifugation to remove cellular debris. To determine infectious virus production, 1 × 10^4^ TZM-bl cells per well were seeded in 96-well cell culture plates and infected in triplicate with the virus-containing supernatants. Viral infectivity was assessed 48-72 h post-infection using the Gal-Screen® kit (Applied Biosystems) according to the manufacturer’s instructions using an Orion microplate luminometer (Berthold). Virus stocks were stored at −80 °C for subsequent analyses.

### HIV-1 sgRNA virus kinetic

To determine the impact of Regnase KO on HIV-1 replication kinetics, CEM-M7-Cas9 and THP-1-Cas9 cells were infected in suspension with HIV-1 virus stocks or VSV-G-pseudotyped HIV-1 stocks, respectively. CEM-M7-Cas9 cells were exposed to virus-containing supernatants for 4 h, whereas THP-1-Cas9 cells were infected overnight. Following infection, cells were washed three times with PBS, resuspended in RPMI 1640 supplemented with 10% FCS, L-glutamine (2 mM), streptomycin (100 μg/mL) and penicillin (100 U/mL) and seeded in triplicate into 96-well plates at a density of 2 × 10⁵ cells per well. Every second day, 150 µL of culture supernatant was collected and stored at −80 °C for subsequent analyses. The sampled volume was replaced with fresh medium, and cells were returned to culture until the next time point. Infectious virus production was quantified by infecting TZM-bl reporter cells with the collected supernatants, followed by measurement of β-galactosidase activity 2-3 days later.

### SDS-PAGE and Western blotting

Cells were lysed in coimmunoprecipitation lysis buffer (150 mM NaCl, 50 mM HEPES, 5 mM EDTA, 0.1% Nonidet P-40, 0.5 mM sodium orthovanadate, 0.5 mM NaF, pH 7.5) supplemented with cOmplete™ EDTA-free protease inhibitor (Roche). Cell lysates were subsequently mixed with Protein Sample Loading Buffer (LI-COR) supplemented with 2-mercaptoethanol. Samples were boiled for 10 min at 95°C and loaded on NuPAGE™ 4–12% Bis-Tris Gels (ThermoFisher Scientific) and transferred onto 0.45 μm pore Immobilon-FL PVDF membranes (Merck Millipore). Membranes were blocked in 5% milk for 1 h and probed using following primary antibodies: Rabbit anti-FLAG (#14793S, Cell signalling), Rat anti-GAPDH (#607902, Biolegend), Mouse anti-HIV-1-p24 (#ARP-3537, NIH AIDS Reagents program) and Mouse anti-HIV-1-env (#ARP-12559, NIH AIDS Reagents program). For detection, blots were stained with dye labelled secondary antibodies (IRDye 680RD goat anti-Rabbit IgG (H + L), #926-68071, LI-COR; IRDye 680RD Goat anti-Mouse IgG (H + L), #926-32210, LI-COR; IRDye 800CW Goat anti-Rat IgG (H + L), #925-32219, LI-COR) and visualized using the LI-COR Odyssey F and Image Studio 6.1 software (LI-COR).

### MLV-GFP expression

To evaluate the effect of Regnase overexpression on MLV gene expression, HEK293T cells were transfected with 750 ng of the pCG N-FLAG plasmid encoding the Regnases sequence and 250 ng of *env*-deficient Moloney MLV eGFP reporter virus DNA^47^. At 72 h post-infection, cells were detached, resuspended in PBS containing 1% FCS and 1% paraformaldehyde (PFA). Cells were analysed by flow cytometry (FACS Canto II) by quantifying the GFP MFI in the BFP+ population (transfected cells).

### RSV-GFP infection

Respiratory Syncytial Virus was grown on HEp-2 cells as described before ^48^. Infectivity of RSV stocks was determined on HEp-2 cells and calculated using the Reed-Muench method ^49^. To evaluate the effect of Regnase overexpression on RSV infection, HEp-2 cells were transfected with 500 ng of the pCG IRES BFP plasmid encoding the Regnases or BFP only. Twenty-four hours after transfection, cells were infected with RSV-GFP at a multiplicity of infection (MOI) of 0.1. At 72 h post-infection, cells were detached, resuspended in PBS containing 1% FCS and fixed in 1% paraformaldehyde (PFA). Cells were analysed by flow cytometry (FACS Canto II) by quantifying the percentage of GFP+ cells (RSV-infected cells) in the BFP+ population (transfected cells).

### HSV-1-GFP infection

HSV-1 GFP stocks were prepared, and their infectivity was determined as described before ^50^. To determine the impact of Regnase overexpression on HSV-1 infection, HEK293T cells (in 96-well plates) were transfected in triplicate with 200 ng of pCG IRES BFP-based Regnase expression plasmids using LT1 transfection reagent according to the manufacturer’s protocol. At 24 h post-transfection, HEK293T cells were infected with HSV-1 GFP reporter virus at 0.01 MOI. GFP fluorescence was assessed using a Cytation imaging multimode reader at 24 h, 48 h, and 72 h post-infection.

### Human coronavirus infection

The method for generating and determining the infectiousness of hCoV-SARS-CoV-2-NL-2020 and OC43 ATCC virus stocks has been described before ^51,52^. To determine the impact of Regnase overexpression on virus infection, HEK293T cells were transfected with 50 ng of ACE2 expression vector and up to 500 ng of Regnase pCG IRES BFP vector or pCG GFP control. After 24 h, cells were infected with 0.1 MOI of SARS-CoV-2 or OC43 virus, and incubated at 37°C (SARS-CoV-2) or 32°C (OC43). The medium was changed the following day, and virus-containing supernatants were harvested 2-3 days post-infection. Viral RNA from the supernatants was isolated using the Qiagen virus RNA isolation kit according to the manufacturer’s recommendations, and the amount of viral RNA was determined using TaqMan 1-step virus master mix and qRT-PCR using the following primers and probes: OC43 forward: agcaaccaggctgatgtcaatacc; OC43 reverse: agcagaccttcctgagccttcaat; OC43 probe: tgacattgtcgatcgggacccaagta; SARS-CoV-2 forward: taatcagac aaggaactgatta; SARS-CoV-2 reverse: cgaaggtgtgacttccatg; SARS-CoV-2 probe: gcaaattgtgcaatttgcgg (synthesized by biomers.net). Copy numbers were calculated using in-house-prepared qRT-PCR standards.

### Cell viability assay

HEK293T cells (0.4 × 10⁶ cells/mL) were seeded in 96-well cell culture plates and transfected with 100 ng of pcDNA4 3xFLAG-Regnase-1-4 expression vector in triplicate. After 48 h, CellTiter-Glo Luminescent Cell Viability Assay (Promega) was performed according to the manufacturer’s recommendations. Luminescence was measured using an Orion microplate luminometer (Berthold).

### Confocal microscopy

HeLa cells were seeded onto glass coverslips in 24-well cell culture plates and transfected with 500 ng of pcDNA4 3xFLAG-Regnase1-4 expression constructs. Cells were treated with leptomycin B (50 nM, 4 h, 37°C) or left untreated prior to fixation. Cells were fixed with 4% paraformaldehyde (PFA) in PBS for 20 min at RT, washed twice with PBS, and permeabilized and blocked in PBS containing 0.5% Triton X-100 and 5% FCS for 1 h. Cells were subsequently incubated with rabbit anti-FLAG (#14793S, Cell signalling) overnight at 4°C. After washing with PBST, cells were incubated with Goat anti-Rabbit Alexa Fluor 647-conjugated secondary antibody (AB_2535813, Invitrogen) and DAPI (Invitrogen) for 2 h at 4°C. Following three washes with PBST, coverslips were rinsed with distilled water, mounted onto microscope slides (ProLong™ Diamond Mounting medium, Cat.#P36965 Invitrogen), and imaged using the Leica DMI 8 fluorescence microscope.

### qRT-PCR

To determine Regnase1-4 expression levels and knock down efficiency, total RNA was extracted using the RNeasy Plus Mini Kit (Qiagen) according to the manufacturer’s instructions. qRT-PCR was performed using the TaqMan™ Fast Virus 1-Step Master Mix (Thermo Fisher Scientific) and a OneStepPlus Real-Time PCR System. All reactions were run in duplicates/triplicates and normalised to GAPDH expression levels or quantified as copy numbers with in-house standard using TaqMan primers/probes from ThermoFisher Scientific: GAPDH (Cat. #4310884E), Regnase-1 (#Hs00962356_m1), Regnase-2 (#Hs01070916_m1), Regnase-3 (#Hs00286919_m1), and Regnase-4 (#Hs01596145_m1).

### IL6-reporter assay

HEK293T cells (0.4 × 10⁶ cells/mL) were seeded in 96-well cell culture plates and transfected the next day with 100 ng of pmirGLO-3’UTR mouse IL6 expression vector and 100 ng of pCG FLAG-Regnase-1-4 expression construct or empty pCG vector as a control. After 48 h, Firefly Luciferase Assay was performed according to the manufacturer’s recommendations (Promega). Luminescence was measured using an Orion microplate luminometer (Berthold).

### HIV-1 LTR reporter assay

To determine the effect of regnases on the untranslated HIV-1 mRNA expressed from HIV-1 LTR promoter containing WT or mutated TAR loop, 20 000 HEK293T cells/well were seeded in 96-well plates and co-transfected in triplicates with firefly luciferase reporter constructs (30 ng) under the control of the HIV-1 NL4-3 LTR and Regnase expression constructs (60 ng) or a control vector (60 ng). At 40 h post-transfection, cells were lysed and firefly luciferase activity was determined using the Luciferase Assay Kit (Promega) according to the manufacturer’s instructions and Orion microplate luminometer (Berthold).

### Phylogenetic and positive selection analysis

Phylogenetic tree showing distance-based relationship inference data of human Regnases were generated using the NGPhylogeny.fr FastME tool (https://ngphylogeny.fr/) and were visualized using iTOL (https://itol.embl.de/)^53^.

MEME (Mixed Effects Model of Evolution), an online tool that employs a mixed-effects maximum-likelihood approach to test the hypothesis that individual sites have been subject to episodic positive or diversifying selection, and FEL (Fixed Effects Likelihood), an online tool that uses a maximum-likelihood (ML) approach to infer nonsynonymous (dN) and synonymous (dS) substitution rates on a per-site basis for a coding alignment and phylogeny to identify selection^54,55^, were used to analyse mammalian Regnase coding obtained from the NCBI gene orthologs database.

### Statistical analysis

Statistical analyses were performed using GraphPad Prism 10 (GraphPad Software). P values were calculated using an unpaired two-tailed Student’s t-test unless otherwise stated. At least three independent experiments were performed unless stated otherwise. Statistical significance is indicated as follows: *p < 0.05, **p < 0.01, and ***p < 0.001. Details of statistical analyses are provided in the figure legends.

## RESULTS

### Regnases inhibit RNA viruses and share common characteristics of antiviral restriction factors

Human Regnases contain a catalytic PIN domain with endoribonuclease activity and a single CCCH-type zinc finger (ZF) that binds RNA (Figure 1A). Despite similar domain organisation, they greatly differ in size (60-120kDa), and their amino acid sequences show a low level of conservation outside of the catalytic PIN domain, suggesting long, distinct evolutionary histories (Figure 1B). To determine if similarly to Regnase-1, Regnases 2-4 also exert antiviral activity, we overexpressed increasing amounts of these Regnases in HEK293T cells together with HIV-1 NL4-3 strain DNA (Figure 1C). At the highest dose, all Regnases inhibited HIV-1 yield by 60-99%. Regnase-1 was the most effective at inhibiting HIV-1 and, even at the lowest tested dose, achieved 93% inhibition. In comparison, inhibition by Regnase-2 reached 53%, and Regnase-3 and -4 overexpression led to a 25% reduction in HIV-1 virus yield at the same DNA concentration. Regnase overexpression also decreased HIV-1 Env RNA levels, although to a lesser extent than the infectious virus yield, suggesting that human Regnases target transcripts encoding Env as well as other genomic regions. Control experiments confirmed that all Regnases were efficiently expressed (Figure 1D, S1B), caused no cytotoxicity in the transfected cells (Figure S1A) and efficiently targeted IL-6 reporter mRNA (Figure S1B) in agreement with published data ^25,28,41,56^. To determine the breadth of the antiviral activity of human Regnases, we compared the effects of their overexpression on diverse RNA and DNA viruses. All human Regnases significantly inhibited the gene expression of Murine Leukemia Virus (MLV), which along with HIV-1 also belongs to the retrovirus family (Figure 1E) as well as infection by diverse clinically relevant single-stranded RNA viruses such as Respiratory Syncytial Virus (RSV) and human coronaviruses OC43, and SARS-CoV-2 (Figures 1F-H). In contrast, none of the human Regnases significantly inhibited infection by the DNA virus Herpes Simplex Virus type 1 (HSV-1) (Figure 1I, S1D). Thus, Regnases show broad antiviral activity against retroviruses and other single-stranded RNA viruses.

**Figure 1.**
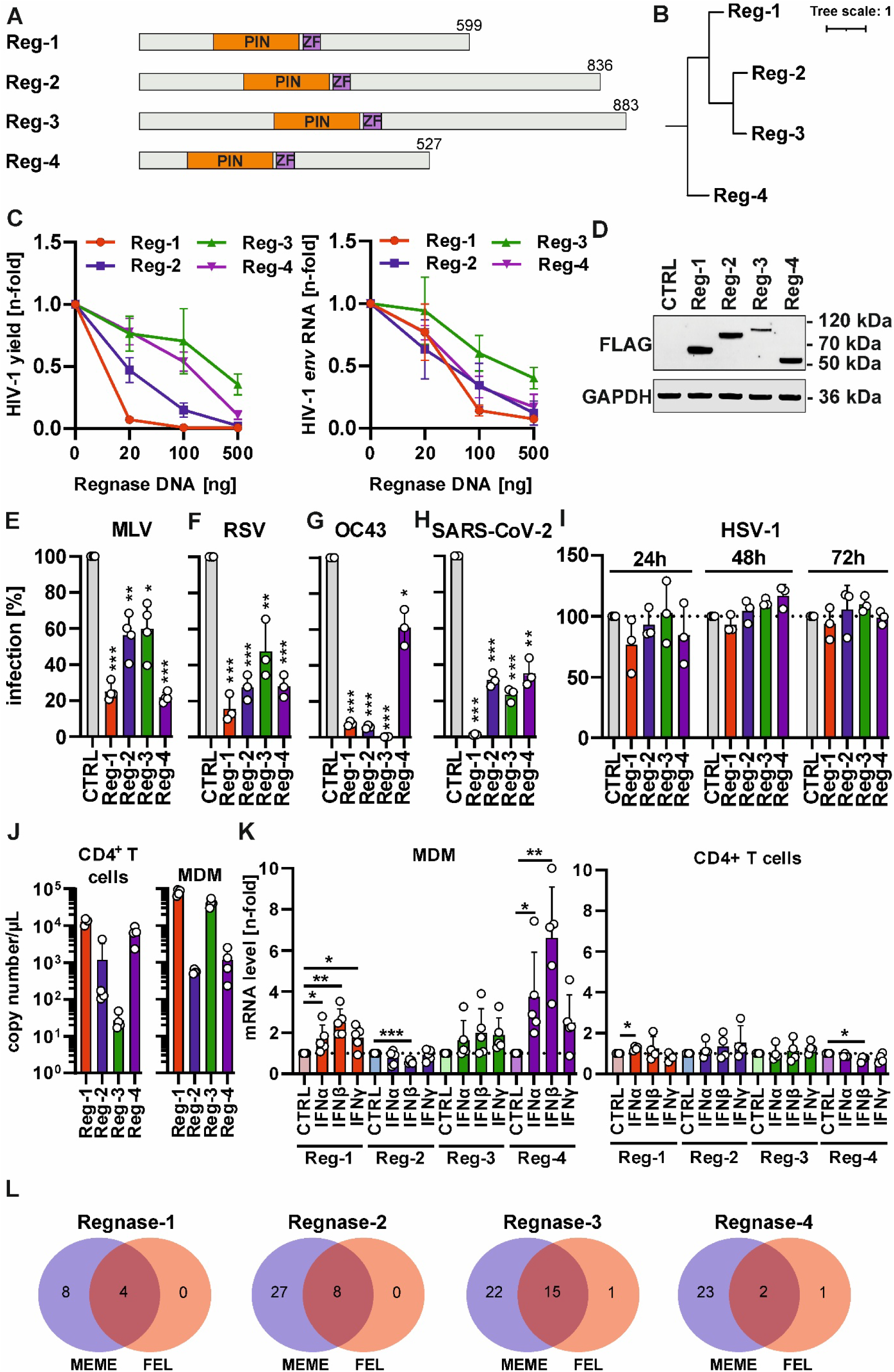
Analysis of the properties of human Regnases as potential antiviral restriction factors. (A) Structural organisation of human Regnases 1-4. The catalytic PilT N-terminus (PIN) endoribonuclease domain and RNA-binding Zinc Finger (ZnF) are shown in orange and purple, respectively. (B) Phylogenetic tree showing the evolutionary relationships among human Regnases, generated from their amino acid sequences. (C) Impact of Regnase overexpression on HIV-1 infectious virus yield and env RNA levels in HIV-1 NL4-3 provirus co-transfected HEK293T cells. Virus yield was determined using TZM-bl HIV-1 reporter cells, while HIV-1 RNA was quantified using qRT-PCR. Mean of n=3 +/− SD, each tested in triplicate. (D) Protein expression levels of FLAG-tagged Regnases in transfected (500 ng DNA) HEK293T cells. GAPDH was used as a loading control. (E) Impact of Regnase overexpression on MLV-GFP provirus expression in co-transfected HEK293T cells. The GFP reporter signal was measured by flow cytometry. (F) Impact of Regnase overexpression on RSV-GFP virus infection in transfected HEp2 cells. The number of GFP+ cells was measured by flow cytometry. (G) Impact of Regnase overexpression on hCoV OC43 (H) and SARS-CoV-2 viral RNA production levels in the supernatant following infection of transfected HEK293T cells co-expressing ACE2. (I) Impact of Regnase overexpression on HSV-1-GFP virus spread in transfected HEK293T cells measured at 24, 48 and 72 h after infection using a Cytation plate imager. (J) RNA expression levels of Regnases in primary human CD4+ T cells and monocyte-derived macrophages (MDM). Mean of 4 individual donors + SD. (K) RNA expression levels of Regnases in primary human MDMs and CD4+ T cells 48 h after stimulation with IFNα (500 U/ml), IFNβ (500 U/ml), IFNγ (200 U/ml). Mean of 4-5 individual donors + SD. (L) Number of sites under positive selection identified by MEME and FEL tools in the Regnase gene coding sequences from mammals. The full list of identified sites is provided in Table S1. Statistical analysis: unpaired Student’s t-test; *, p < 0.05; **, p < 0.01; ***, p<0.001.

Since HIV-1 is the best-described viral target of Regnase-1^17^, we sought to determine whether other Regnases are expressed in primary target cells of HIV-1 *in vivo*, CD4+ T cells and macrophages^57^. To do this, we isolated both cell types from human blood and quantified the mRNA copy numbers of the Regnases using qRT-PCR (Figure 1J). Regnase-1 was expressed in both cell types, Regnase-3 was poorly expressed in primary T cells but highly expressed in monocyte-derived macrophages (MDMs), whereas Regnase-4 showed the opposite expression pattern. Meanwhile, Regnase-2 was expressed at low levels in both cell types, which is consistent with previous reports ^25^. Since the expression of antiviral factors is often upregulated by interferons (IFNs), we also tested IFN-inducibility of human Regnases in primary cells (Figure 1K). Analysis of Regnase mRNA expression in the presence of type I and II interferons (IFNs) revealed that Regnase-1 and Regnase-4 are further induced by IFNs, however, they only respond well to IFN treatment in MDMs, but not in primary CD4+ T cells. In contrast, Regnases-2 and -3 were not induced by IFN treatment, with Regnase-2 and Regnase-4 levels showing even a minor significant downmodulation in MDMs and CD4+ T cells, respectively. The lack of meaningful induction in CD4+ T cells was not due to the timing of stimulation, as kinetic measurements of Regnase mRNA levels between 2-72 h post-stimulation did not reveal more than 2-fold differences in expression at any of the tested time points (Figure S1E). Finally, since restriction factors often evolve under positive selection, we compared the signatures of evolutionary selection in mammalian Regnase genes (Figure 1L). The analysis revealed that all members of the Regnase family contain multiple sites showing signs of positive evolutionary selection (Figure 1L). Thus, human Regnases share several features of classical antiviral restriction factors^1^: they show broad antiviral activity, have evolved under positive selection, and Regnase-1 and -4 are upregulated by IFNs.

### Differential sensitivity of HIV-1 and HIV-2 and related simian lentiviruses to human Regnases

Host restriction factors co-evolve together with viral pathogens in an evolutionary arms race. This is particularly evident in the case of HIV and SIVs, which are highly diverse due to high error rates of the viral reverse transcriptase. This high genomic diversity enables evasion of many known restriction factors^1^. Since the HIV-1 NL4-3 strain has been extensively cell-line passaged and may have lost some of its original immune-evasive properties^58,59^, we first decided to compare the impact of Regnases on a set of patient-derived isolates of HIV-1 subtype B, which is dominant in North America, Western and Central Europe, and Australia (Figure 2A). All strains were efficiently inhibited by Regnase-1 and to a lesser degree by Regnase-4, with no major differences between HIV-1 NL4-3 and the primary strains. In the case of Regnase-2 and Regnase-3, the degree of inhibition varied, with the NL4-3 and CH058 strains being less restricted than the CH077 and RHGA strains. Since the Regnase-2 gene (*ZC3H12B*) is located on the X chromosome and its expression levels are higher in females^60^, we hypothesized that viruses isolated from female patients might have evolved increased resistance to this endoribonuclease. To test this, we compared primary HIV-1 subtype C isolates obtained from a female (CH167) and male (CH042) patient (Figure 2B). However, there were no statistically significant differences between these two strains for any of the tested Regnases. To further confirm these results, we compared Regnase-2 sensitivity of an extended set of 10 chronic subtype C strains isolated from male (5) and female (5) patients (Figure 2C). This extended analysis found no evidence that patient sex influences Regnase-2 sensitivity of HIV-1 isolates.

**Figure 2.**
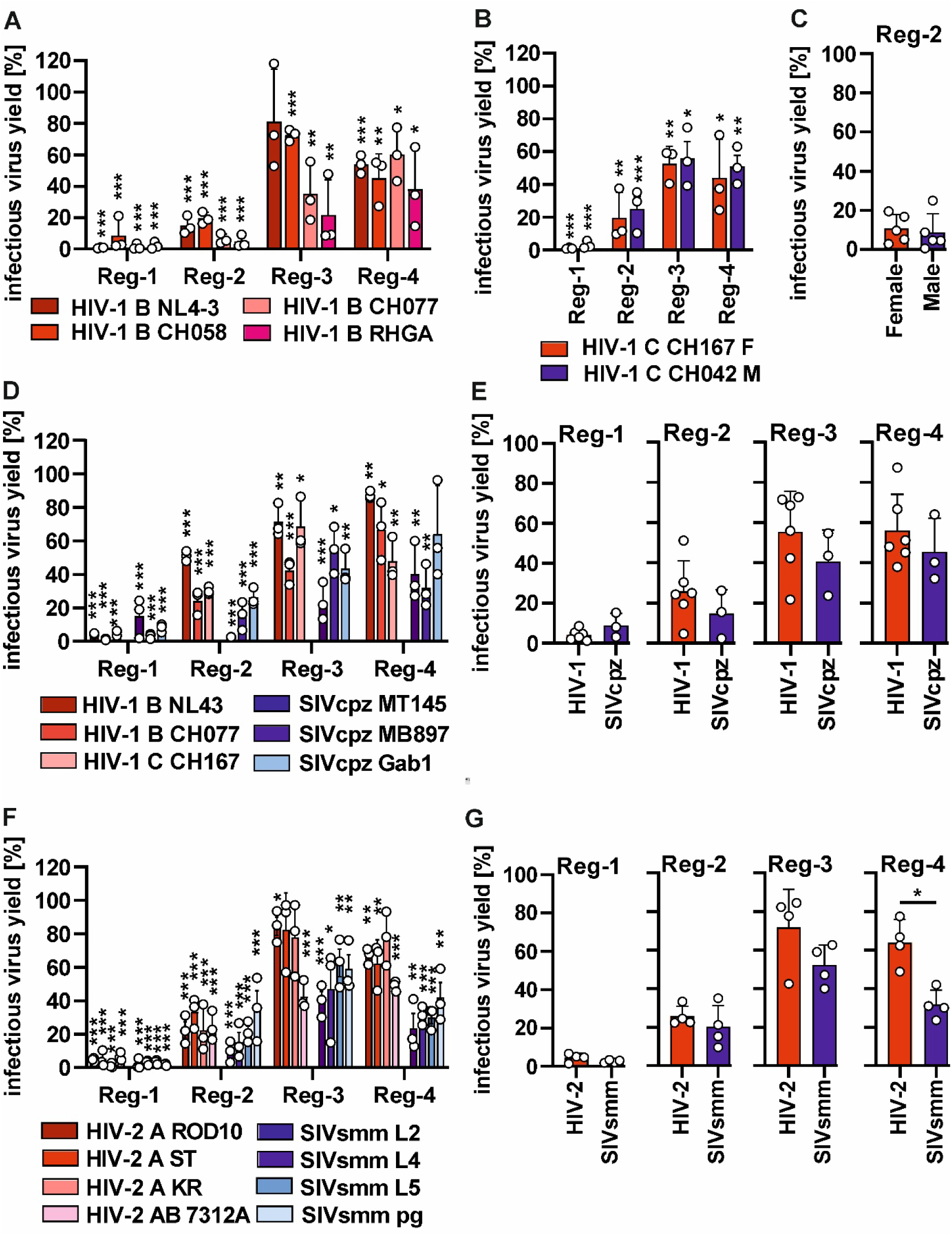
Differences in viral sensitivity to human Regnases. (A) Impact of Regnase overexpression on HIV-1 infectious virus yield in indicated HIV-1 subtype B provirus co-transfected HEK293T cells. Statistical comparison shown was calculated for each virus transfected with regnase or empty vector control (to which the values were normalised as 100%) (B, C) Impact of Regnase overexpression on HIV-1 infectious virus yield of indicated HIV-1 proviruses isolated from a male (M) and female (F) patients in transfected HEK293T cells and (D) Impact of Regnase overexpression on HIV-1 and SIVcpz infectious virus yield in co-transfected HEK293T cells and (E) a group comparison between HIV-1 (red) and SIVcpz (blue) viruses. (F) Impact of Regnase overexpression on HIV-2 and SIVsmm infectious virus yield from co-transfected HEK293T cells and (G) group comparison between HIV-2 (red) and SIVsmm (blue) viruses. Infectious virus yield was determined using TZM-bl HIV-1 reporter cells. Statistical analysis: panels D and F: Mann-Whitney test; all other panels: unpaired Student’s t-test. *, p < 0.05; **, p < 0.01; ***, p<0.001.

To determine whether human Regnases might have contributed to the interspecies barrier during viral transmission from chimpanzees to humans and host-specific adaptation, we compared the Regnase sensitivity of HIV-1 and its animal precursor SIVcpz (Figure 2D). On average, SIVcpz strains were slightly more sensitive to human Regnases-2-4, however, the differences between the groups were not statistically significant (Figure 2E). In contrast, HIV-2 and its animal precursor SIVsmm showed a significant 2-fold difference in sensitivity to Regnase-4 (Figure 2F-G). Importantly, these differences did not stem from lower infectivity of simian viruses as there was no significant correlation between Regnase sensitivity and absolute virus yield (Figure S2). This suggests that while the antiviral activity of human Regnases is very broad, HIV-2 have evolved greater resistance to human Regnase-4, and this endoribonuclease could have contributed to the interspecies barrier that limits the successful transmission of monkey-infecting SIVs to humans.

### Endogenous human Regnases inhibit HIV-1 infection in T cells and macrophages

Since overexpression assays likely involve higher-than-physiological levels of antiviral proteins, it was important to confirm that endogenously expressed Regnases also exert antiviral activity. To test this, we took advantage of the recently developed “traitor virus” system, which induces a gene knockout (KO) in HIV-1-infected cells ^37,61^. The sgRNA encoded by a cassette inserted into the replication-competent virus enables amplification of antiviral phenotypes over multiple replication cycles (Figure 3A). Since this system requires Cas9-expressing cell lines, we first compared Regnase mRNA expression in two established HIV-1 cell line-based infection models: CEM-M7 and THP-1 cells stably expressing Cas9 (Figure 3B and 3C). As expected, Regnase-2 expression was low in both cell lines, consistent with levels observed in primary cells (Figure 1J), whereas the other family members were expressed at intermediate (Regnase-1 and -3) or high levels (Regnase-4). We selected 2 sgRNAs targeting each Regnase gene and determined their targeting efficiency in Cas9-electroporated CEM-M7 cells. Due to low Regnase protein expression levels and the lack of suitable antibodies, we were unable to quantify the knockout at the protein level; however, DNA analysis of these cells revealed that all sgRNAs, except Regnase-1 sgRNA 1, effectively targeted their corresponding genes (Figure 3D). The sgRNAs had no impact on the infectivity of the traitor virus stock produced in HEK293T cells (Figure 3E). Furthermore, in the absence of Cas9 expression in CEM-M7 cells, these viruses had no advantage over the non-targeting (NT) sgRNA control (Figure 3F). However, in the presence of Cas9, viruses encoding sgRNAs targeting ZAP (positive control) as well as human Regnases-1, -2 and -4 produced 1.5- to 3.5-fold higher levels of infectious HIV-1 than the non-targeting control (NT) (Figure 3G, SsA). In the case of Regnase-3, only one of the tested sgRNAs showed a modest advantage (1.8-fold). We also measured the replication of HIV-1 sgRNA viruses in the THP-1 Cas9 monocyte-like cells (Figure 3H). While Regnase-4 KO was associated with the highest replication advantage (up to 3.2-fold), surpassing that of ZAP, Regnase-1 (∼2-fold), Regnase-2 and -3 had little to no effect (1-1.8-fold). To further verify these results in primary human cells, we performed Regnase knockdown using siRNA in primary MDMs. Knockdown of Regnase-1 and Regnase-3 was highly efficient, while Regnase-4 mRNA levels were reduced by ∼50% (Figure 3I). Regnase-2 expression levels were reduced in 3 of 4 donors, with large variability likely due to low absolute Regnase-2 expression levels (Figure 1J). Surprisingly, despite the low transcript levels, the knockdown of Regnase-2 resulted in the most pronounced enhancement of HIV-1 infection (Figure 3J), followed by Regnase-3 and Regnase-4. Surprisingly, despite its high antiviral activity in overexpression assays and high knockdown efficiency of Regnase-1, viral infection was unaffected by Regnase-1 knockdown in MDMs. Since none of the tested infection models showed high levels of Regnase-2, we investigated the antiviral phenotype of Regnase-2 in the glioblastoma-derived cell line U373 MAGI (Figure S3B). This cell line was poorly permissive to HIV-1 sgRNA viruses; therefore, we pseudotyped the virus stocks with VSV-G, which boosted viral infection (Figure S3C) and measured the impact of sgRNAs on a single round of virus infection (Figure S3D). While sgRNAs targeting ZAP, Regnase-3, and Regnase-4 led to a significant 3-5-fold increase in infectious virus production, neither Regnase-1 nor Regnase-2 had a pronounced antiviral effect in this system. Thus, human Regnase-1, -2, and -3 inhibit HIV-1 infection at endogenous levels in a cell-type-dependent manner, while Regnase-4 inhibits the virus in all tested cell types.

**Figure 3.**
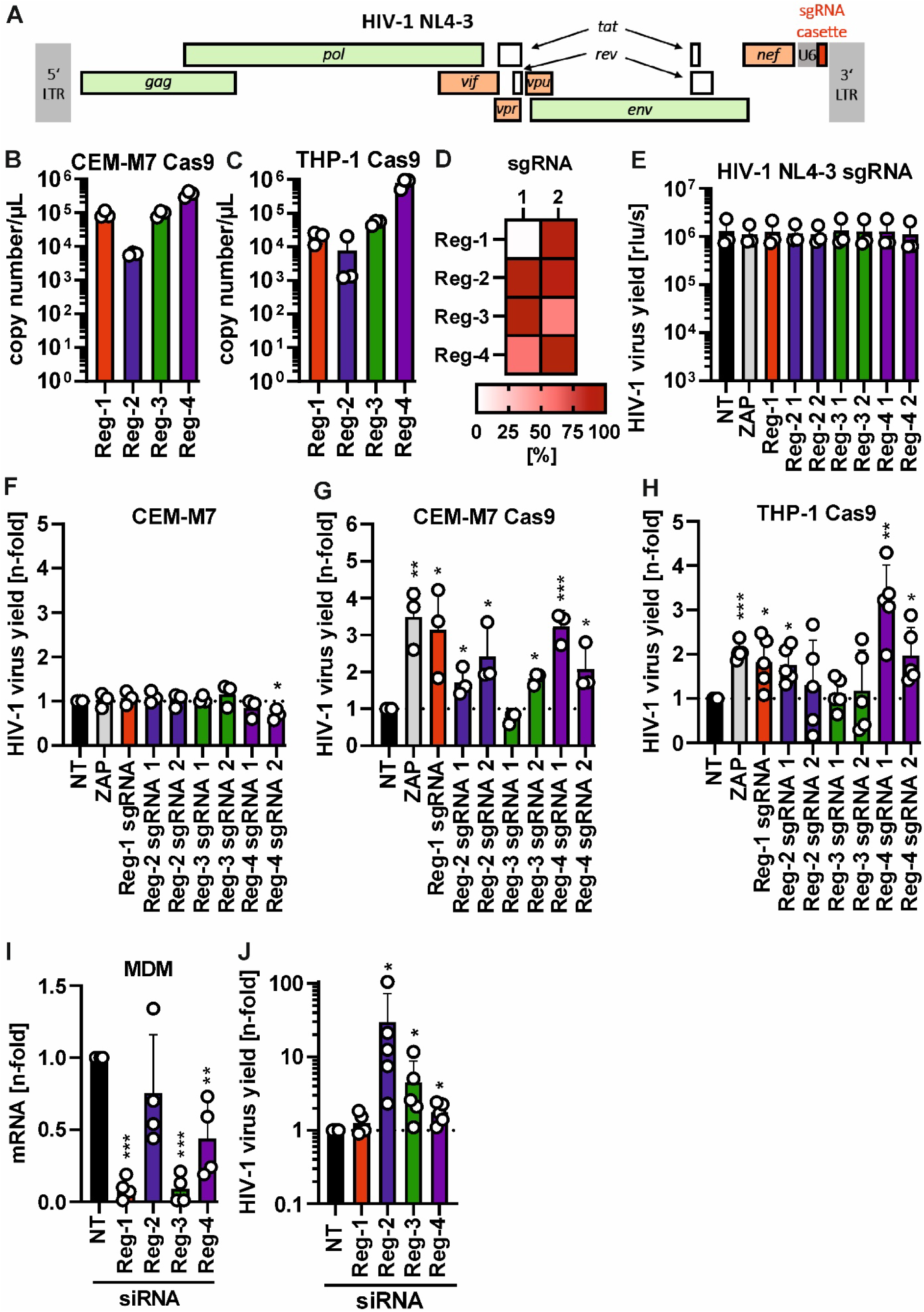
Antiviral activity of endogenous human Regnases against HIV-1. (A) Schematic of the modified HIV-1 NL4-3 viral genome containing the sgRNA cassette for infection-conditional host gene knockout in Cas9-expressing cells. (B) Expression levels of Regnase mRNA in CEM-M7 Cas9 and (C) THP-1 Cas9 cells. (D) Efficiency of Regnase gene targeting (% gene disruption) by selected sgRNAs in CEM-M7 cells as compared to non-targeting (NT) sgRNA. (E) Mean infectious virus yield of HIV-1 NL4-3 viruses containing sgRNAs targeting ZAP or human Regnases generated by transfection into HEK293T cells. Mean infectious virus yield from infected CEM-M7 not expressing Cas9 (F) and (G) expressing Cas9 (mean of days 8-18 of the kinetic) and (H) THP-1 Cas9 cells (mean of days 3-12 of the kinetic) compared to NT control. Mean of n=3-5 +SD. (I) Efficiency and (J) enhancing effect of siRNA knockdown of ZAP and Regnases in primary human monocyte-derived macrophages (MDMs) from 5 different donors. Statistical analysis: unpaired Student’s t-test. *, p < 0.05; **, p < 0.01; ***, p<0.001.

### Antiviral activity of Regnases requires intact catalytic, RNA-recognition, and dimerisation sites

The PIN and CCCH zinc-finger domains are the most conserved parts of the Regnase family of proteins (Figure 4A). They mediate three properties of Regnases: catalytic endoribonuclease activity, dimerisation (both PIN domain; orange), RNA binding/recognition (Zinc Finger; purple), all of which are suggested to be important for cellular RNA targeting by Regnase-1 ^11,43,62^. To determine whether the targeting of viral RNA by Regnases also involves these sites, we mutated the first catalytic site in each Regnase and tested their antiviral activity in overexpression assays against HIV-1 NL4-3 (Figure 4B) and CH077 strains (Figure 4C). This mutation led to a significant, albeit incomplete, loss of antiviral activity across all Regnases. To determine whether residual antiviral activity was mediated by the other aspartic acid residues that form the catalytic core, we mutated all four sites combined and tested its impact on HIV-1 CH077 strain (Figure 4D, Figure S4A). These mutants behaved similarly to the single-site mutants, which were also efficiently expressed (Figure 4E), suggesting that 20-40% of the observed inhibition might arise from cleavage-independent properties of Regnases, such as RNA binding. To test this hypothesis, we mutated the CCCH residues to AAAA to disrupt RNA recognition/binding (Figure 4F-G). The ZnF mutants of Regnase-3 and -4 were completely inactive, while the mutants of Regnase-1 and -2 retained significant antiviral activity against the HIV-1 CH077 virus (Figure 4G). However, when both catalytic and ZnF residues were mutated in Regnase-1 and -2, it led to a complete loss of antiviral activity (Figure 4H-I). Interestingly, the putative dimerisation site of the PIN domain (PR motif) was absolutely required for the antiviral activity of Regnase-1, -2 and -3 (Figure 4J-K), but not for Regnase-4. This effect could also not be explained by the expression levels of Regnase-4 mutants (Figure 4L), and structural dimer predictions produced by AlphaFold did support dimerisation with a similar level of confidence to other Regnases (Figure S4B). Therefore, it remains to be determined whether Regnase-4 acts as a monomer, dimerises at other sites or heterodimerises with endogenously expressed Regnases in HEK293T cells (Figure S4C). Overall, these results show that antiviral activity of Regnases is functionally dependent on their catalytic, RNA-recognition and dimerisation properties.

**Figure 4.**
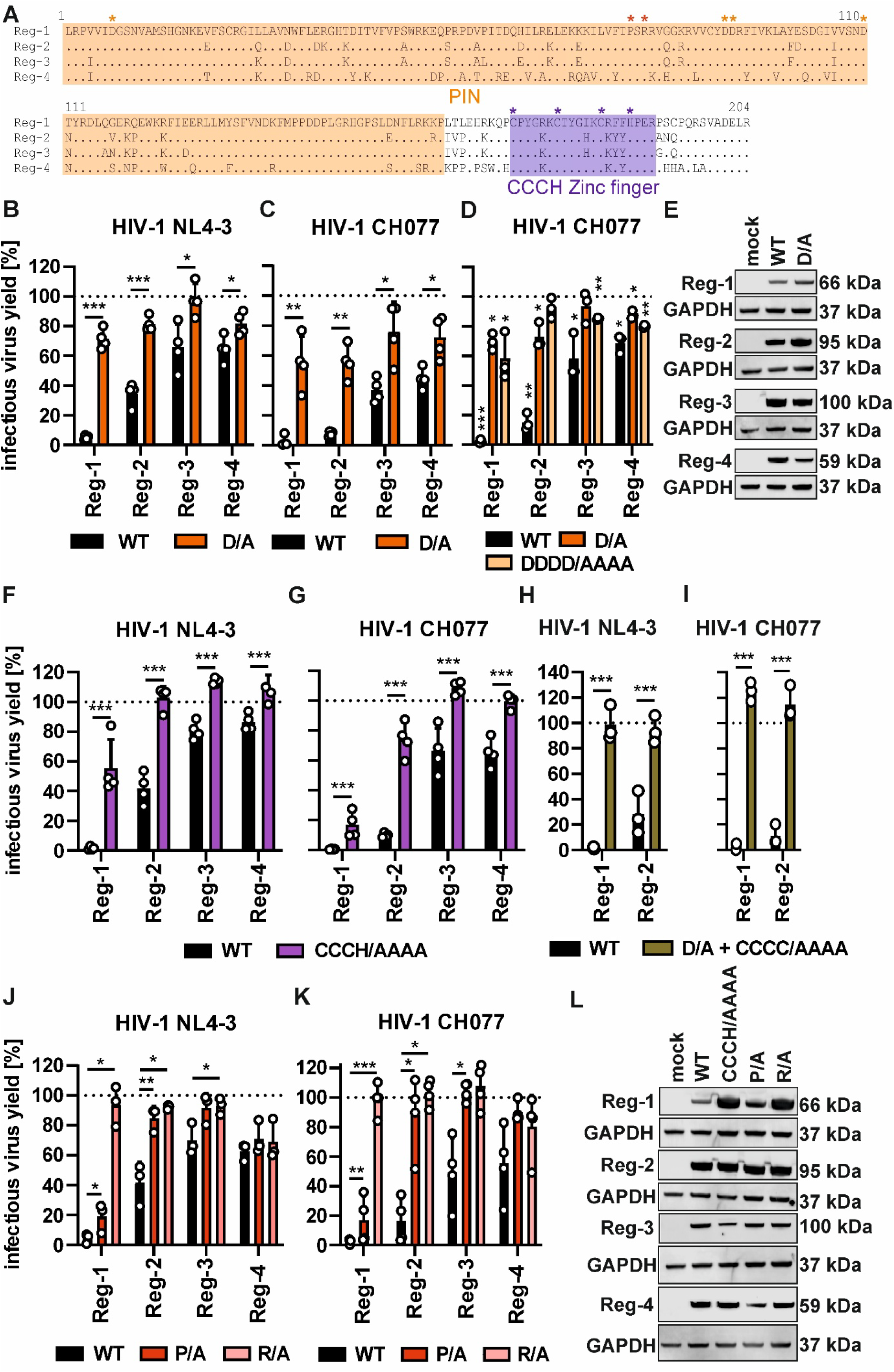
Molecular determinants of antiviral activity of human Regnases. (A) Alignment of the PIN (orange) and CCCH Zinc-finger (purple) domains of the human Regnases. The exact location of the four catalytic D residues forming the catalytic core, ZnF CCCH residues binding RNA and putative dimerisation sites is indicated with stars. (B) Impact of wild type (WT) or catalytic mutant (D/A) Regnase overexpression on HIV-1 NL4-3 or (C) CH077 infectious virus yield in co-transfected HEK293T cells. (D) Effect of quadruple catalytic site mutation in Regnases on HIV-1 CH077 infectious virus yield in co-transfected HEK293T cells. (E) Western blot showing protein expression levels of FLAG-tagged Regnases and the D/A mutants. (F) Impact of wild type (WT) or ZnF mutant (CCCH/AAAA) Regnase overexpression on HIV-1 NL4-3 or (G) CH077 infectious virus yield in co-transfected HEK293T cells. (H) Impact of the combined catalytic site and ZnF mutation of Regnase-1 and -2 on HIV-1 NL4-3 and (I) CH077 infectious virus yield. (J) Impact of wild-type (WT) or dimerisation mutant (P/A and R/A) Regnase overexpression on HIV-1 NL4-3 or (K) CH077 infectious virus yield in co-transfected HEK293T cells. (L) Western blot showing protein expression levels of FLAG-tagged WT Regnases and the CCCH/AAAA and P/A and R/A mutants. Mean of n=3-4 +SD. Statistical analysis: unpaired Student’s t-test. *, p < 0.05; **, p < 0.01; ***, p<0.001.

### Regnases-1, -3 and -4 shuttle between the cytoplasm and nucleus

As the antiviral potency of the Regnase family members differs, we decided to further determine which properties are linked to their antiviral phenotypes. Since HIV-1 produces its RNA in the nucleus and later exports it into the cytoplasm, where viral protein expression and genomic RNA packaging occur, we hypothesised that localisation to these compartments is required for viral RNA targeting. To determine the subcellular localisation of Regnases, we overexpressed FLAG-tagged proteins in HeLa cells. The localisation of all Regnases was diffuse and cytoplasmic (Figure 5A), with no indication of specific nuclear localisation. However, upon treatment with the nuclear CRM1 export inhibitor leptomycin B, Regnases-1, -3 and -4 localised predominantly in the nucleus, suggesting that they are nuclear shuttling proteins (Figure 5B). Meanwhile, Regnase-2 did not change its localisation under treatment and remained in the cytoplasm, suggesting that nuclear transport is not required for its antiviral activity. The differences between Regnase-2 and other family members were confirmed by quantification of FLAG and DAPI colocalization in multiple cells (Figure 5C). Thus, even though at the steady state all Regnases localise to the cytoplasm and likely target viral RNA there, Regnases-1, 3 and -4 might also shuttle in and out of the nucleus, where HIV-1 RNA synthesis takes place.

**Figure 5.**
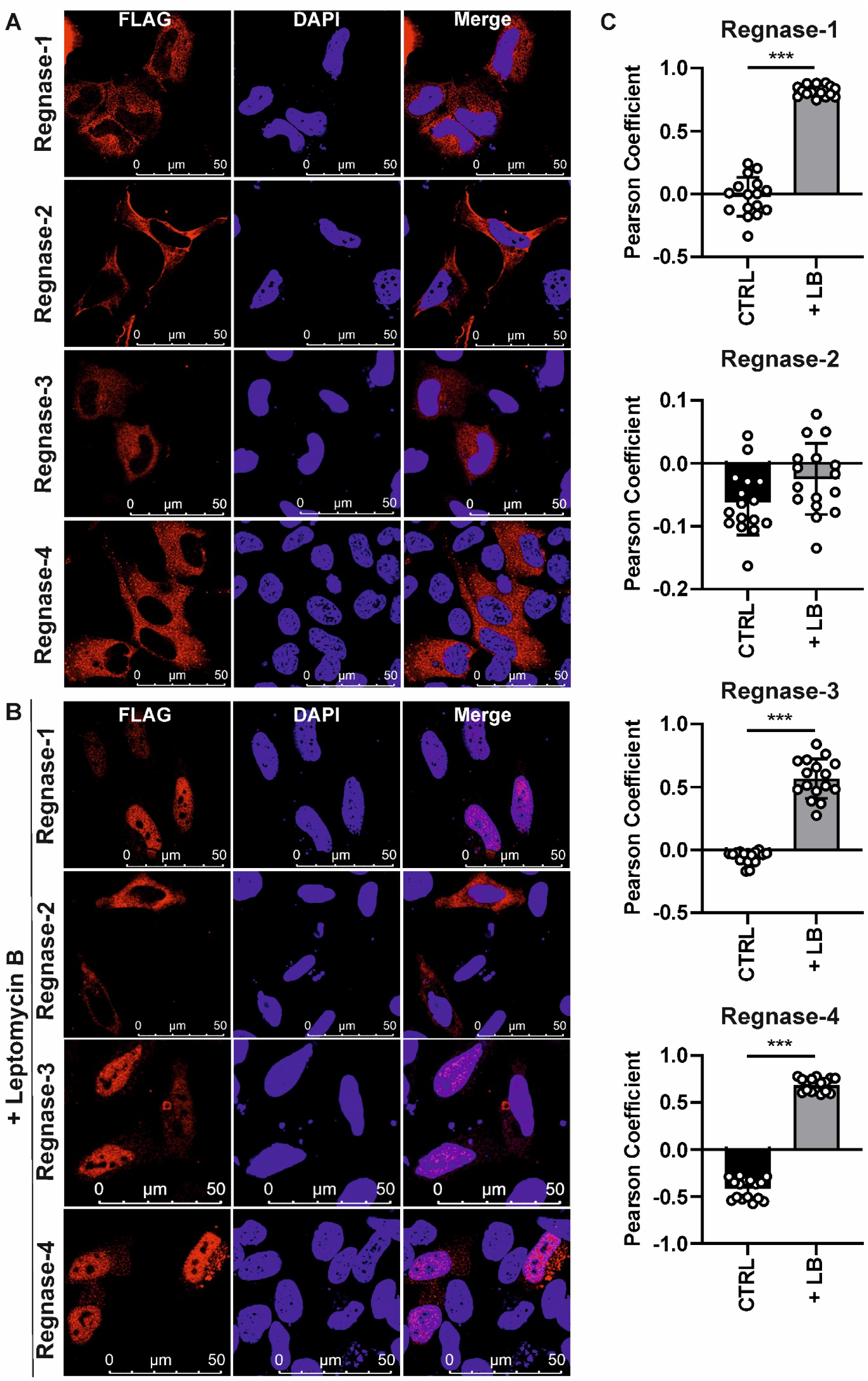
Subcellular localisation of human Regnases. (A) Confocal microscopy images of FLAG-tagged human Regnases overexpressed in human HeLa cells (B) and following 4 h treatment with nuclear export inhibitor Leptomycin B (LB). DAPI served as a nuclear marker. (C) Quantification of Regnase and DAPI colocalization signal (Pearson coefficient) in the absence and presence of LB treatment. Each dot represents one analysed cell; bars indicate mean ± SD. ***, p<0.001.

### Regnases have different HIV-1 RNA specificity

To dissect if differences in catalytic activity or RNA recognition are primarily responsible for differential activity of Regnases, we swapped the N and C-terminal halves of the most and least active Regnases (−1 and −3; Figure 6A). These chimeric Regnases contained the N-terminus including the catalytic PIN domain of one member of the family and C-terminus containing the RNA-binding domain (ZnF) of another one, allowing the dissection of the effects of RNA cleavage and recognition on antiviral activity. The N-terminal (including the catalytic domain) swap had a minor effect on antiviral activity against HIV-1, compared to C-terminal swap (containing the ZnF domain) (Figure 6B). This suggested that the differences in the antiviral potencies of human Regnases might result from differences in ZnF-mediated viral RNA recognition or binding.

**Figure 6.**
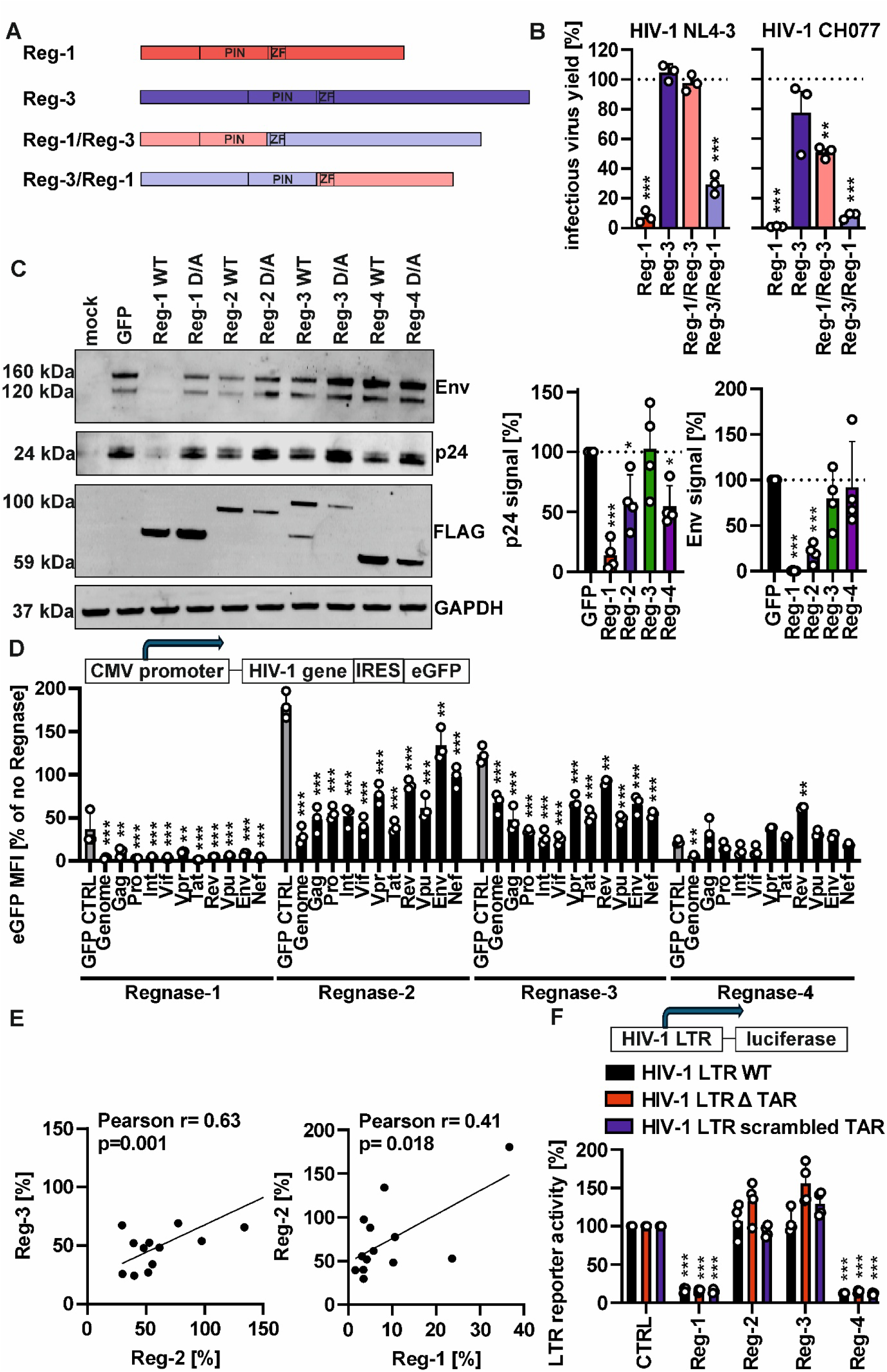
Comparison of target specificity of human Regnases. (A) Schematic of generated Regnase-1 and Regnase-3 chimeric proteins and (B) Impact of wild type (WT) or chimeric Regnase construct overexpression on HIV-1 NL4-3 or CH077 infectious virus yield in co-transfected HEK293T cells. (C) Western blot showing the expression of HIV-1 Env and Gag (p24) proteins in the presence of FLAG-tagged WT and catalytic mutant Regnase expression in co-transfected HEK293T cells. Right: quantification of p24 and Env signal in the presence of WT Regnases. GAPDH served as a loading and normalisation control. (D) Upper panel: schematic of IRES eGFP reporter constructs encoding individual HIV-1 proteins. Lower panel: Impact of Regnase overexpression on different IRES eGFP reporter activity, normalised to empty control vector for each viral protein (100%). Statistical analysis compares the individual viral proteins to GFP control (grey bars) for each Regnase. (E) Correlation between HIV-1 protein IRES eGFP reporter activity in the presence of Regnase-2 and -3, or -1 and -2 overexpression. Pearson coefficient and p values are indicated. (F) Upper panel: schematic of HIV-1 LTR luciferase reporter construct insert. Lower panel: Impact of Regnase overexpression on HIV-1 LTR activity of WT, TAR deleted and TAR scrambled reporters. Mean of n=3-4 +SD. Statistical analysis: unpaired Student’s t-test. *, p < 0.05; **, p < 0.01; ***, p<0.001.

To determine what type of viral RNA products were targeted by Regnases, we first examined HIV-1 CH077 Gag (p24) and Env expression levels in the presence of individual Regnases (Figure 6C). While Gag is expressed from the unspliced viral RNA, Env-encoding transcripts are singly spliced. Regnase-1 and -2 decreased the expression levels of both viral proteins, Regnase-3 had no significant effect while Regnase-4 decreased Gag levels but had no impact on Env. To investigate which other HIV-1 protein-coding regions are also efficiently targeted by Regnases, we took advantage of an HIV-1 protein expression construct library containing a reporter IRES eGFP element that allows tracking of viral mRNA levels through GFP measurements (Figure 6D). Since Regnase-1 activity in overexpression assays was much more potent than that of other family members, we used a two-fold lower dose of this Regnase. To exclude off-target effects resulting from non-specific targeting, we also measured the effect of Regnases on the GFP reporter expression and used this to determine statistically significant differences (Figure 6D, S5C). The expression vector encoding HIV-1 reverse transcriptase (RT) was poorly expressed and therefore excluded from further analysis. The expression of viral reporter genes in Regnase-expressing cells varied, with all types being significantly more suppressed by Regnase-1, -2 and -3 compared to the GFP control. Meanwhile, the whole HIV-1 genome but none of the individual genes was significantly suppressed by Regnase-4, due to strong inhibition of the GFP reporter itself. Statistical analyses revealed a significant correlation between the targeting specificity of Regnase-2 and -3 and, to a lesser extent, Regnase-1 and -2 (Figure 6E). In contrast, there was no significant correlation between Regnase-4 and any other Regnase family member, suggesting that its effects in this assay are less specific.

Apart from protein-coding regions, HIV-1 RNA also contains untranslated regions (UTRs) that fulfil regulatory functions and can be targeted by RNA-degrading proteins. Since Regnase-1 has been reported to target a hairpin structure in the UTR of the IL-6 mRNA transcript, we investigated whether the HIV-1 UTR, and more specifically the highly conserved Trans-activation response element (TAR) loop, is also targeted by these endoribonucleases. To do this, we employed a reporter construct that expresses the firefly luciferase under the control of HIV-1 long terminal repeat (LTR) containing TAR (Figure 6F). To control for possible non-specific targeting of the luciferase gene by Regnases, we used a luciferase expression construct under the control of CMV promoter to normalise the data (Figure S5D). While Regnase-1 and -4 both strongly inhibited reporter activity, Regnase-2 and -3 had no significant specific effects. Furthermore, mutagenesis of the TAR loop, either through complete deletion or sequence randomisation (scrambling), had no impact on targeting by Regnase-1 and -4, suggesting that other UTR elements mediate viral RNA recognition and targeting.

Thus, the high antiviral potency of Regnase-1 is not linked to its superior catalytic activity but rather to a broad specificity that encompasses both translated and untranslated regions of HIV-1 genome. Similarly, Regnase-4, which also inhibited HIV-1 infection at endogenous expression levels in T cells (Figure 3G), suppressed the HIV-1 LTR independently of the TAR sequence, which is consistent with targeting of other untranslated LTR sequence elements. Meanwhile, Regnase-2 and Regnase-3 seem to predominantly target translated viral RNA regions.

## DISCUSSION

This study provides a systematic comparison of all four human Regnases as restriction factors and establishes that antiviral activity is a conserved property of the human Regnase family. While previous studies have primarily focused on the ability of Regnase-1 to suppress viral replication by degrading viral RNA ^14,15,17–19^, the antiviral potential of Regnases-2, -3, and -4 against different RNA viruses remained largely unexplored. This is likely due to an early report showing that Japanese Encephalitis Virus and Dengue Virus are not efficiently targeted by overexpressed Regnases 2-4^19^. Here, we demonstrate that all four family members inhibit diverse RNA viruses, including HIV-1, HIV-2, MLV, RSV and coronaviruses SARS-CoV-2 and OC43, but not the DNA virus HSV-1. Endogenous depletion experiments further demonstrated that Regnases contribute to the intrinsic restriction of HIV-1 in human T cells, monocyte- and glia-derived cell lines, as well as primary macrophages. Mechanistically, by focusing on HIV-1 as a well-established virus-infection model for studying antiviral RNA-binding proteins^32,63,64^, we show that efficient antiviral activity requires intact catalytic, ZnF, and, for most family members, dimerisation motifs, while comparative analyses reveal substantial differences in antiviral potency, cell-type specific expression, subcellular localisation, and viral RNA specificity among individual Regnases. Together, these findings identify the human Regnase family as antiviral RNases with distinct specificities that allow complementary control of viral RNA.

Several characteristics identified in this study support the classification of Regnases as antiviral restriction factors. Classical restriction factors often directly interfere with viral replication by recognising conserved viral features, are constitutively expressed and further upregulated by interferons, frequently evolve under positive selection, and are evaded or antagonized by highly successful viruses^1^. Human Regnases fulfil all (Regnase-1 and -4) or most (Regnases-2 and -3) of these criteria. All family members suppress viral RNA expression during overexpression; endogenous Regnases significantly restrict HIV-1 replication to a similar degree as ZAP; Regnase-1 and Regnase-4 are induced by type I interferons in primary macrophages; and all four genes exhibit signatures of positive selection across mammals. HIV-1 was previously found to evade Regnase-1 by replicating in activated T cells, where Regnase-1 is cleaved and inactivated by paracaspase MALT1^17^. Furthermore, in this study, we found that Regnase-4 is more antiviral against SIVsmm than against HIV-2, suggesting that the latter has at least partially evolved to evade or counteract this restriction factor during its human adaptation. These observations are consistent with evolutionary pressure and virus-host interactions acting on Regnases. Signatures of positive selection have been reported as a defining feature of many established antiviral proteins, including APOBEC3 proteins, tetherin, and ZAP ^1,63,65–67^, strengthening the argument for classifying Regnase family enzymes as novel antiviral restriction factors.

A crucial observation emerging from this work is that despite conserved structural architecture, antiviral activity is not uniform across the Regnase family or in different cell types. Regnase-1 consistently displayed the strongest antiviral activity in overexpression assays, consistent with previous reports demonstrating inhibition of HIV-1, HCV, HBV, and coxsackievirus B3 ^15–19,22^. It also showed one of the highest efficiencies in restricting HIV-1 infection in CEM-M7 Cas9 cells, but had only a marginal effect in monocytic THP-1 Cas9 cells and no effect in primary cell-derived macrophages. While no previous studies have examined the antiviral function of Regnase-1 in primary human macrophages, the T cell data are consistent with published data showing that Regnase-1 efficiently restricts HIV-1 infection in resting T cells^17^. This suggests that the endogenous antiviral function of Regnase-1 might be cell-type-dependent, possibly due to post-translational mechanisms that regulate Regnase-1 protein levels, such as paracaspase- or caspase-mediated cleavage^68–70^. In contrast to Regnase-1, Regnase-4 exhibited potent endogenous antiviral activity in both T cells and monocytic cell lines while showing rather modest performance in overexpression assays. Unexpectedly, despite its relatively low mRNA expression levels, Regnase-2 showed strong antiviral phenotypes in primary human macrophages but not other cell types, whereas Regnase-3 displayed intermediate behaviour that depended on both the virus strain and cellular context. These observations challenge the findings of a previous overexpression study that compared the antiviral activity of Regnases against flaviviruses and suggested that only Regnase-1 has antiviral activity^19^. While it is possible that the mechanism and potency of Regnase targeting of these two viruses differ from that of HIV-1, our study clearly shows that antiviral properties are determined not only by intrinsic enzymatic activity but also by expression levels and recognition of specific motifs in viral RNA in both translated and untranslated regions. Therefore, endogenous knockdown and knockout experiments provide the most accurate estimate of physiological contribution, while overexpression experiments are valuable for mechanistic studies. Regnase mRNA expression differed substantially between primary CD4+ T cells and macrophages, and only Regnase-1 and Regnase-4 responded robustly to interferon stimulation in macrophages. These findings suggest that different family members contribute to IFN-mediated intrinsic immunity in distinct cell types. Such specialisation is consistent with previous studies implicating Regnase-3 predominantly in myeloid cells and Regnase-4 in immune regulation within lymphoid compartments. The relatively weak mRNA induction observed in primary T cells also emphasises that antiviral activity cannot simply be inferred from interferon responsiveness, as post-transcriptional regulation and protein stability also influence the function of human regnases ^68–70^. It remains to be established what the contribution of the individual Regnases to viral restriction is in other relevant HIV-1 infection models, such as lymphoid tissues and primary CD4+ T cells, and how immune activation affects Regnase-mediated restriction of HIV-1 and other RNA viruses.

Our data indicate that Regnases preferentially target RNA viruses. All tested RNA viruses were susceptible to inhibition, whereas HSV-1 replication was completely unaffected despite comparable experimental conditions. Interestingly, it has been shown that Regnase-1 can target other DNA viruses, such as KSHV and EBV ^20,21^. Thus, our data suggest that HSV-1 antagonises or evades the activity of human Regnases, or that Regnases specifically target HSV-1 RNA transcripts, which have no effect on viral spread *in vitro*. The exact mechanism of HSV-1’s resistance to Regnases requires further study.

An important unanswered question is whether viruses have evolved mechanisms to evade or antagonize Regnase-mediated restriction. Numerous antiviral restriction factors are counteracted by dedicated viral antagonists or even exploited by viruses ^1,71^. Although no viral protein has yet been shown to directly inhibit Regnases, HIV-1 infection mainly replicates in activated T cells, which induces MALT1-dependent cleavage of Regnase-1, thereby allowing the virus to evade the antiviral activity^68^. Other viruses may similarly evade Regnase restriction by modulating Regnase expression, subcellular localization, post-translational modification or protein stability. Alternatively, viral RNAs may evolve structural features that reduce recognition by Regnases while preserving essential functions. The differential sensitivity observed among HIV-2 and SIV strains is consistent with the possibility that lentiviruses may have already acquired partial resistance to Regnase-4 during human adaptation. Identifying viral determinants that modulate Regnase activity and defining the RNA features recognised by individual Regnases will therefore be important for understanding the evolutionary interplay between these cellular RNases and RNA viruses.

Previous structural and biochemical studies established that catalytic activity and dimerisation mediated by the PIN-domain, as well as RNA binding mediated through the CCCH zinc finger, contribute to the degradation of cellular transcripts by Regnase-1 ^25,28,43^. Our results demonstrate that these same molecular determinants are also required for efficient antiviral activity. Loss of catalytic activity substantially impaired restriction by all family members, confirming that RNA cleavage represents the dominant antiviral mechanism. Nevertheless, catalytic mutants retained some residual antiviral activity, particularly in Regnase-1 and Regnase-2, suggesting that RNA recognition itself may interfere with viral gene expression independently of endonucleolytic cleavage, for example, through translational repression of bound RNA. This possibility is supported by the complete loss of activity following simultaneous disruption of catalytic and ZnF sites in Regnase-1 and -2, which could not be achieved by selective mutation of either motif. Similar cleavage-independent antiviral mechanisms have been proposed for other RNA-binding restriction factors, including ZAP, which recognises viral RNA and recruits additional RNA-degrading proteins to degrade it ^5,72,73^. Whether the optimal antiviral activity of Regnases requires additional RNA sensors or components of the cellular RNA degradation machinery remains to be investigated in future studies.

The significant role of dimerisation sites in the antiviral activity of Regnases also provides a new mechanistic insight into their mechanism of action. Disruption of the putative PIN-domain dimerisation interface abolished the activity of Regnases-1, -2 and -3 but not Regnase-4. Structural studies suggested that Regnase-1 functions as a functional dimer during RNA degradation^43^, and our data indicate that this mechanism may extend to additional family members. The apparent independence of Regnase-4 from this interface may reflect an alternative mode of oligomerisation, functional heterodimerisation with PIN domains of other family members, or differences in structural organisation. Future structural analyses comparing all four family members should help explain these functional differences and may reveal additional determinants underlying their distinct antiviral activities.

One of the most intriguing findings of this study is that differences in antiviral potency appear to arise primarily from substrate recognition rather than catalytic activity. The exchange of domains between Regnase-1 and Regnase-3 demonstrated that antiviral activity largely depends on the C-terminal region containing CCCH ZnF rather than the N-terminal region with the catalytic PIN domain. Consistent with this interpretation, reporter assays revealed distinct viral RNA targeting profiles among family members. Regnase-1 and Regnase-4 targeted a wide range of substrates, including genes of non-viral origin, whereas Regnases-2 and -3 exhibited more selective patterns of inhibition and a significant correlation in their substrate preferences. These findings suggest that diversification within the Regnase family has primarily occurred at the level of RNA recognition while preserving a conserved catalytic mechanism. Such functional specialization would allow different family members to collectively recognize a wider repertoire of structured viral and cellular RNAs than could be achieved by a single enzyme.

Our work argues against a single specific genomic region, such as the highly structured UTR, being the primary determinant of Regnase-mediated recognition. Although the lentiviral TAR loop represents an attractive candidate because of its highly structured RNA architecture and functional similarities to the IL-6 stem-loop previously identified as a Regnase-1 substrate, deletion or scrambling of HIV-1 TAR did not abolish its targeting by Regnase-1 and -4. Instead, our findings suggest that other elements within the viral untranslated regions or coding sequences are targeted by Regnases. HIV-1 contains numerous conserved RNA secondary structures that regulate reverse transcription, splicing, nuclear export and genome packaging, many of which remain plausible candidates for Regnase recognition. Comprehensive transcriptome-wide mapping of Regnase binding sites using approaches such as CLIP-seq will therefore be required to define the structural features underlying their substrate specificity.

Our work has several limitations. Much of the mechanistic characterisation relies on transient overexpression, which may not fully recapitulate endogenous protein abundance or regulation. Although endogenous knockdown and conditional knockout experiments confirmed physiological antiviral activity, these analyses were performed for HIV-1, and it remains to be determined whether endogenous Regnases similarly restrict other RNA viruses. Furthermore, while our data strongly support targeting of viral RNA, the precise RNA structures recognised by individual Regnases remain unidentified. Finally, given the well-established immunoregulatory functions of Regnases ^11,13^, dissecting direct antiviral activity from the effects on cellular gene expression will require future studies.

In summary, this study establishes the human Regnase family as an important component of intrinsic antiviral immunity. Rather than functioning as redundant RNases, individual Regnases exhibit distinct expression patterns, regulatory properties and RNA substrate preferences that together broaden the spectrum of antiviral defence. These findings substantially expand the biological role of Regnases beyond regulation of inflammatory mRNA stability and suggest that evolutionary diversification of RNA recognition has enabled this family to participate in host defence against diverse RNA viruses. Understanding how Regnases discriminate between viral and cellular RNAs, how viruses evade their activity, and how these pathways intersect with innate immune signalling will provide important insights into both antiviral immunity and RNA biology, and may identify new opportunities for therapeutic manipulation of host antiviral responses.

## ACKNOWLEDGEMENTS

We thank Dré van der Merwe, Regina Burger, Martha Mayer, Jana-Romana Fischer, Daniela Krnavek, Birgit Ott, and Nicola Schrott for laboratory assistance. We thank Dr Alexandre Laliberté, Dr Caterina Prelli-Bozzo, Dr Fabian Zech and Eszter Matiz for technical advice and for sharing resources. This manuscript has been written with the help of artificial intelligence (AI). All AI edits and information were critically reviewed by the authors prior to publication.

This work was supported by the European Union’s Horizon 2020 research and innovation programme under Marie Sklodowska-Curie (grant agreement No. 101062524 to D.K.), Else-Kröner Fresenius Stiftung (2022_EKEA.47 to D.K.) and DFG (KM 5/3-1 to D.K. and K.S.). F.K. was supported by an ERC Advanced grant (Traitor viruses) and the DFG (KI 548/21-1). D.S. was funded by the Heisenberg Program (grant ID: SA 2676/3-1) of the German Research Foundation (DFG). K.S. was supported by the DFG (SP 1600/13-1, CRC1506-INST 40/657-3).

## DATA AVAILABILITY STATEMENT

The primary data underlying this article as well as generated resources will be shared on a reasonable request to the corresponding author.

## Notes

### Competing Interest Statement

The authors have declared no competing interest.

### Summary of Updates

New figures and additional authors have been added.

## REFERENCES

1. Kluge, S. F., Sauter, D. & Kirchhoff, F. SnapShot: antiviral restriction factors. Cell 163, 774–774.e1 (2015).

2. Nchioua, R., Bosso, M., Kmiec, D. & Kirchhoff, F. Cellular Factors Targeting HIV-1 Transcription and Viral RNA Transcripts. Viruses 12, 495 (2020).

3. Kmiec, D. & Kirchhoff, F. Antiviral factors and their counteraction by HIV-1: many uncovered and more to be discovered. Journal of Molecular Cell Biology mjae005 (2024) doi:10.1093/jmcb/mjae005.

4. Yamasoba, D. et al. N4BP1 restricts HIV-1 and its inactivation by MALT1 promotes viral reactivation. Nat Microbiol 4, 1532–1544 (2019).

5. Ficarelli, M. et al. KHNYN is essential for the zinc finger antiviral protein (ZAP) to restrict HIV-1 containing clustered CpG dinucleotides. Elife 8, e46767 (2019).

6. Clemens, M. J. & Williams, B. R. G. Inhibition of cell-free protein synthesis by pppA2′ p5′ A2′ p5′ A: a novel oligonucleotide synthesized by interferon-treated L cell extracts. Cell 13, 565–572 (1978).

7. Kerr, I. M. & Brown, R. E. pppA2’p5’A2’p5’A: an inhibitor of protein synthesis synthesized with an enzyme fraction from interferon-treated cells. Proceedings of the National Academy of Sciences 75, 256–260 (1978).

8. Hovanessian, A. G., Brown, R. E. & Kerr, I. M. Synthesis of low molecular weight inhibitor of protein synthesis with enzyme from interferon-treated cells. Nature 268, 537–540 (1977).

9. Baglioni, C., Minks, M. A. & Maroney, P. A. Interferon action may be mediated by activation of a nuclease by pppA2′p5′A2′p5′A. Nature 273, 684–687 (1978).

10. Li, X.-L., Blackford, J. A. & Hassel, B. A. RNase L Mediates the Antiviral Effect of Interferon through a Selective Reduction in Viral RNA during Encephalomyocarditis Virus Infection. J Virol 72, 2752–2759 (1998).

11. Muzio, L., Ferrati, D., Colombo, E. & Molinaro, C. The Regnase pathway: a core axis in immune regulation and inflammatory disease. Front. Immunol. 17, 1756491 (2026).

12. Uehata, T. & Takeuchi, O. Post-transcriptional regulation of immunological responses by Regnase-1-related RNases. Int Immunol 33, 859–865 (2021).

13. Fischer, M., Weinberger, T. & Schulz, C. The immunomodulatory role of Regnase family RNA-binding proteins. RNA Biol 17, 1721–1726 (2020).

14. Li, Y. et al. MCPIP1 reduces HBV-RNA by targeting its epsilon structure. Sci Rep 10, 20763 (2020).

15. Lin, R.-J. et al. MCPIP1 Suppresses Hepatitis & Virus Replication and Negatively Regulates Virus-Induced Proinflammatory Cytokine Responses. J Immunol 193, 4159–4168 (2014).

16. Li, M. et al. MCPIP1 inhibits coxsackievirus B3 replication by targeting viral RNA and negatively regulates virus-induced inflammation. Med Microbiol Immunol 207, 27–38 (2018).

17. Liu, S. et al. MCPIP1 restricts HIV infection and is rapidly degraded in activated CD4+ T cells. Proceedings of the National Academy of Sciences 110, 19083–19088 (2013).

18. Li, M. et al. MCPIP1 inhibits Hepatitis B virus replication by destabilizing viral RNA and negatively regulates the virus-induced innate inflammatory responses. Antiviral Research 174, 104705 (2020).

19. Lin, R.-J. et al. MCPIP1 ribonuclease exhibits broad-spectrum antiviral effects through viral RNA binding and degradation. Nucleic Acids Res 41, 3314–3326 (2013).

20. Kook, I. & Ziegelbauer, J. M. Monocyte chemoattractant protein-induced protein 1 directly degrades viral miRNAs with a specific motif and inhibits KSHV infection. Nucleic Acids Res 49, 4456–4471 (2021).

21. Happel, C., Ramalingam, D. & Ziegelbauer, J. M. Virus-Mediated Alterations in miRNA Factors and Degradation of Viral miRNAs by MCPIP1. PLOS Biology 14, e2000998 (2016).

22. Li, H. & Wang, T. T. MCPIP1/regnase-I inhibits simian immunodeficiency virus and is not counteracted by Vpx. Journal of General Virology 97, 1693–1698 (2016).

23. Lichawska-Cieslar, A. et al. MCPIP1 functions as a safeguard of early embryonic development. Sci Rep 13, 16944 (2023).

24. Liang, J. et al. A Novel CCCH-Zinc Finger Protein Family Regulates Proinflammatory Activation of Macrophages*. Journal of Biological Chemistry 283, 6337–6346 (2008).

25. Wawro, M. et al. ZC3H12B/MCPIP2, a new active member of the ZC3H12 family. RNA 25, 840–856 (2019).

26. Long, J. et al. The role of ZC3H12D-regulated TLR4-NF-κB pathway in LPS-induced pro-inflammatory microglial activation. Neurosci Lett 832, 137800 (2024).

27. Zhang, H. et al. ZC3H12D attenuated inflammation responses by reducing mRNA stability of proinflammatory genes. Mol Immunol 67, 206–212 (2015).

28. Wawro, M., Kochan, J., Krzanik, S., Jura, J. & Kasza, A. Intact NYN/PIN-Like Domain is Crucial for the Degradation of Inflammation-Related Transcripts by ZC3H12D. Journal of Cellular Biochemistry 118, 487–498 (2017).

29. Liu, B. et al. The RNase MCPIP3 promotes skin inflammation by orchestrating myeloid cytokine response. Nat Commun 12, 4105 (2021).

30. Liu, L. et al. Zc3h12c inhibits vascular inflammation by repressing NF-κB activation and pro-inflammatory gene expression in endothelial cells. Biochem J 451, 55–60 (2013).

31. Villenave, R. et al. Induction and Antagonism of Antiviral Responses in Respiratory Syncytial Virus-Infected Pediatric Airway Epithelium. Journal of Virology 89, 12309–12318 (2015).

32. Kmiec, D. et al. HIV-2 evades restriction by ZAP through adaptations in the U3 LTR region despite increased CpG levels. Nucleic Acids Res 53, gkaf826 (2025).

33. Kmiec, D. et al. CpG Frequency in the 5′ Third of the env Gene Determines Sensitivity of Primary HIV-1 Strains to the Zinc-Finger Antiviral Protein. mBio 11, 10.1128/mbio.02903-19 (2020).

34. Parrish, N. F. et al. Phenotypic properties of transmitted founder HIV-1. Proceedings of the National Academy of Sciences 110, 6626–6633 (2013).

35. Fenton-May, A. E. et al. Relative resistance of HIV-1 founder viruses to control by interferon-alpha. Retrovirology 10, 146 (2013).

36. Parrish, N. F. et al. Transmitted/Founder and Chronic Subtype & HIV-1 Use CD4 and CCR5 Receptors with Equal Efficiency and Are Not Inhibited by Blocking the Integrin α4β7. PLOS Pathogens 8, e1002686 (2012).

37. Prelli Bozzo, C., et al. Replication competent HIV-guided CRISPR screen identifies antiviral factors including targets of the accessory protein Nef. Nat Commun 15, 3813 (2024).

38. Olari, L.-R. et al. α-Synuclein fibrils enhance HIV-1 infection of human T cells, macrophages and microglia. Nature Communications 16, 813 (2025).

39. Platt, E. J., Wehrly, K., Kuhmann, S. E., Chesebro, B. & Kabat, D. Effects of CCR5 and CD4 cell surface concentrations on infections by macrophagetropic isolates of human immunodeficiency virus type 1. J Virol 72, 2855–2864 (1998).

40. Derdeyn, C. A. et al. Sensitivity of human immunodeficiency virus type 1 to the fusion inhibitor T-20 is modulated by coreceptor specificity defined by the V3 loop of gp120. J Virol 74, 8358–8367 (2000).

41. Wawro, M. et al. Molecular Mechanisms of ZC3H12C/Reg-3 Biological Activity and Its Involvement in Psoriasis Pathology. Int J Mol Sci 22, 7311 (2021).

42. Wawro, M., Kochan, J. & Kasza, A. The perplexities of the ZC3H12A self-mRNA regulation. Acta Biochim Pol 63, 411–415 (2016).

43. Yokogawa, M. et al. Structural basis for the regulation of enzymatic activity of Regnase-1 by domain-domain interactions. Sci Rep 6, 22324 (2016).

44. Leoni, C., Bataclan, M., Ito-Kureha, T., Heissmeyer, V. & Monticelli, S. The mRNA methyltransferase Mettl3 modulates cytokine mRNA stability and limits functional responses in mast cells. Nat Commun 14, 3862 (2023).

45. Hotter, D. et al. IFI16 Targets the Transcription Factor Sp1 to Suppress HIV-1 Transcription and Latency Reactivation. Cell Host & Microbe 25, 858–872.e13 (2019).

46. Brinkman, E. K., Chen, T., Amendola, M. & van Steensel, B. Easy quantitative assessment of genome editing by sequence trace decomposition. Nucleic Acids Res 42, e168 (2014).

47. Erdemci-Evin, S. et al. A Variety of Mouse PYHIN Proteins Restrict Murine and Human Retroviruses. Viruses 16, (2024).

48. Hunszinger, V. et al. Respiratory viruses activate autophagy via the IFN-STAT1/STAT5B-SOCS1 axis. 2025.10.28.685013 Preprint at 10.1101/2025.10.28.685013 (2025).

49. Reed, L. J. & Muench, H. A SIMPLE METHOD OF ESTIMATING FIFTY PER CENT ENDPOINTS12. American Journal of Epidemiology 27, 493–497 (1938).

50. Freisem, D. et al. Herpes simplex virus infection promotes ALS pathology through ICP0-mediated PML body disruption. 2026.03.27.714707 Preprint at 10.64898/2026.03.27.714707 (2026).

51. Nchioua, R. et al. SARS-CoV-2 Is Restricted by Zinc Finger Antiviral Protein despite Preadaptation to the Low-CpG Environment in Humans. mBio 11, e01930–20 (2020).

52. Xie, Q., et al. Endogenous IFITMs boost SARS-coronavirus 1 and 2 replication whereas overexpression inhibits infection by relocalizing ACE2. iScience 26, 106395 (2023).

53. Lemoine, F., et al. NGPhylogeny.fr: new generation phylogenetic services for non-specialists. Nucleic Acids Research 47, W260–W265 (2019).

54. Kosakovsky Pond, S. L. & Frost, S. D. W. Not So Different After All: A Comparison of Methods for Detecting Amino Acid Sites Under Selection. Mol Biol Evol 22, 1208–1222 (2005).

55. Murrell, B. et al. Detecting Individual Sites Subject to Episodic Diversifying Selection. PLOS Genetics 8, e1002764 (2012).

56. Yaku, A. et al. Regnase-1 Prevents Pulmonary Arterial Hypertension Through mRNA Degradation of Interleukin-6 and Platelet-Derived Growth Factor in Alveolar Macrophages. Circulation 146, 1006–1022 (2022).

57. Pan, X., Baldauf, H.-M., Keppler, O. T. & Fackler, O. T. Restrictions to HIV-1 replication in resting CD4+ T lymphocytes. Cell Res 23, 876–885 (2013).

58. Jacob, R. A. et al. The HIV-1 accessory protein Nef increases surface expression of the checkpoint receptor Tim-3 in infected CD4+ T cells. J Biol Chem 297, 101042 (2021).

59. Kmiec, D. et al. Vpu-Mediated Counteraction of Tetherin Is a Major Determinant of HIV-1 Interferon Resistance. mBio 7, 10.1128/mbio.00934-16 (2016).

60. Oliva, M. et al. The impact of sex on gene expression across human tissues. Science 369, eaba3066 (2020).

61. Xie, Q. et al. Replication-competent SIVcpz CRISPR screen identifies barriers to successful cross-species transmission. bioRxiv 2025.10.27.684817 (2025) doi:10.1101/2025.10.27.684817.

62. Mino, T. & Takeuchi, O. Regnase-1-related endoribonucleases in health and immunological diseases. Immunol Rev 304, 97–110 (2021).

63. Nchioua, R. et al. APOBEC3F Constitutes a Barrier to Successful Cross-Species Transmission of Simian Immunodeficiency Virus SIVsmm to Humans. Journal of Virology 95, 10.1128/jvi.00808-21 (2021).

64. Kmiec, D., Lista, M. J., Ficarelli, M., Swanson, C. M. & Neil, S. J. D. S-farnesylation is essential for antiviral activity of the long ZAP isoform against RNA viruses with diverse replication strategies. PLOS Pathogens 17, e1009726 (2021).

65. McLaren, P. J. et al. Identification of potential HIV restriction factors by combining evolutionary genomic signatures with functional analyses. Retrovirology 12, 41 (2015).

66. Uriu, K., Kosugi, Y., Ito, J. & Sato, K. The Battle between Retroviruses and APOBEC3 Genes: Its Past and Present. Viruses 13, 124 (2021).

67. Sauter, D. et al. Tetherin-Driven Adaptation of Vpu and Nef Function and the Evolution of Pandemic and Nonpandemic HIV-1 Strains. Cell Host & Microbe 6, 409–421 (2009).

68. Uehata, T. et al. Malt1-Induced Cleavage of Regnase-1 in CD4+ Helper T Cells Regulates Immune Activation. Cell 153, 1036–1049 (2013).

69. Jeltsch, K. M. et al. Cleavage of roquin and regnase-1 by the paracaspase MALT1 releases their cooperatively repressed targets to promote TH17 differentiation. Nat Immunol 15, 1079–1089 (2014).

70. Suk, F.-M. et al. MCPIP1 Enhances TNF-α-Mediated Apoptosis through Downregulation of the NF-κB/cFLIP Axis. Biology (Basel) 10, 655 (2021).

71. Prelli Bozzo, C., Kmiec, D. & Kirchhoff, F. When good turns bad: how viruses exploit innate immunity factors. Curr Opin Virol 52, 60–67 (2022).

72. Zhu, Y., Wang, X., Goff, S. P. & Gao, G. Translational repression precedes and is required for ZAP-mediated mRNA decay. EMBO J 31, 4236–4246 (2012).

73. Lista, M. J. et al. A Nuclear Export Signal in KHNYN Required for Its Antiviral Activity Evolved as ZAP Emerged in Tetrapods. Journal of Virology 97, e00872–22 (2023).

