## Supplementary Figures for "Human Regnases are antiviral restriction factors targeting viral RNA"

**Figure S1. Expression levels of human Regnases.** Related to Figure 1. (A) Impact of GFP (CTRL) or Regnase overexpression (500 ng) on cell viability measured by Cell-Titer Glo cell viability assay. DMSO (30%) served as a positive control. Mean of n=3 + SD. (B) Impact of GFP (CTRL) or Regnase overexpression (500 ng) on luciferase expression from the IL-6 UTR reporter. Mean of n=4 + SD. (C) Uncropped western blots from Figure 1. Protein expression of FLAG-tagged human Regnases was analyzed in HEK293T cells transiently transfected with 500 ng of the indicated expression plasmids. GAPDH served as a loading control. (D) Representative images of HSV-1-GFP infection in Regnase-overexpressing HEK293T cells. HEK293T cells were transiently transfected with BFP-tagged Regnase expression constructs or a BFP control vector and after 24 h infected with HSV-1-GFP (MOI 0.01). Representative GFP fluorescence images were acquired at 72 h post-infection using a Cytation imaging reader. (E) Primary human $CD4^+$ T cells were left untreated or stimulated with IFNα (500 U/mL), IFNβ (500 U/mL), or IFNγ (200 U/mL) for 2–72 h. Cells were then harvested, and Regnase mRNA expression levels were quantified by qRT-PCR. Shown are the relative mRNA expression levels, expressed as fold change compared with mock-treated cells. Data represent the mean ± SD of four individual donors.

**Figure S2. Correlations between Regnase sensitivity and absolute infectivity of tested HIV-1 strains.** Related to Figure 2.

**Figure S3. Extended characterization of Regnase-mediated restriction of HIV-1 replication.** (A) Replication kinetics of HIV-1 NL4-3 viruses containing sgRNAs targeting ZAP or human Regnases in CEM-M7 Cas9 cells. Infectious virus production was monitored over time (days 2–20) and quantified as infectious virus yield (RLU/s). Individual replication curves are shown. (B) Expression levels of endogenous Regnase-1, Regnase-2, Regnase-3, and Regnase-4 mRNA in U373-MAGI cells, determined by RT-qPCR and expressed as copy number/µL. (C) Replication of non-pseudotyped and VSV-G-pseudotyped HIV-1 NL4-3 sgRNA constructs (non-targeting [NT] or ZAP-targeting) in U373-MAGI cells. (D) Effect of sgRNA-mediated targeting of ZAP and human Regnases on HIV-1 replication in U373-MAGI cells at day 6 post infection. Infectious virus yield was determined following infection with HIV-1 NL4-3 viruses containing the respective sgRNA cassettes and compared with the non-targeting (NT) sgRNA control. Data are presented as mean ± SD from three independent experiments.

**Figure S4. HEK293T expression and AlphaFold2-Multimer prediction of Regnase-1–4 PIN domain homodimers.** (A) Western blot showing the expression of HIV-1 Env and Gag (p24) proteins in the presence of FLAG-tagged WT and 1x (D/A) or 4x catalytic mutant (DDDD/AAAA) Regnase expression in co-transfected HEK293T cells. GAPDH served as a loading control. (B) Predicted PIN domain homodimer structures of Regnase-1, Regnase-2, Regnase-3, and Regnase-4 generated with AlphaFold2-Multimer. The highest-ranked (rank 1) model is shown for each protein. PIN domain chain A is colored blue and chain B gray. Conserved proline residues are highlighted in yellow (Regnase-1 P212, Regnase-2 P267, Regnase-3 P322, Regnase-4 P166), conserved arginine residues in green (Regnase-1 R214, Regnase-2 R269, Regnase-3 R324, Regnase-4 R168), and catalytic aspartate residues in magenta (Regnase-1 D141, D225, D226, D244; Regnase-2 D196, D280, D281, D299; Regnase-3 D251, D335, D336, D354; Regnase-4 D95, D179, D180, D198). Model confidence metrics were: Regnase-1 (pLDDT = 96.2, pTM = 0.907, ipTM = 0.870), Regnase-2 (pLDDT = 95.6, pTM = 0.873, ipTM = 0.807), Regnase-3 (pLDDT = 95.3, pTM = 0.878, ipTM = 0.823), and Regnase-4 (pLDDT = 95.8, pTM = 0.897, ipTM = 0.843). (C) RNA expression levels of Regnase-1, Regnase-2, Regnase-3, and Regnase-4 in HEK293T cells, determined by RT-qPCR and expressed as copy number/µL. Data are presented as mean ± SD from three independent experiments.

**Figure S5. Expression analysis of Regnase variants and their effects on HIV-1 NL4-3 protein expression.** (A) Uncropped western blots showing the expression of HIV-1 NL4-3 Env and Gag (p24) proteins in the presence of FLAG-tagged wild-type (WT) and catalytic mutant Regnase proteins in co-transfected HEK293T cells. GAPDH served as a loading control. (B) Western blot showing the expression of FLAG-tagged Regnase-1 and Regnase-3 wild-type proteins, as well as Regnase-1/Regnase-3 and Regnase-3/Regnase-1 chimeric proteins, in transfected HEK293T cells. GAPDH served as a loading control. (C) Impact of Regnase overexpression on GFP expression. Mean of n=3 +SD. (D) Impact of Regnase overexpression on luciferase expression from a CMV and HIV-1 NL4-3 LTR promoters. Mean of n=3 +SD. Statistical analysis: unpaired Student's t-test. *, $p < 0.05$; **, $p < 0.01$; ***, $p<0.001$.

**Table S1. Sites under positive selection in mammalian Regnases identified using the MEME and FEL tools.**

Grabe et al., 2026
Figure S1

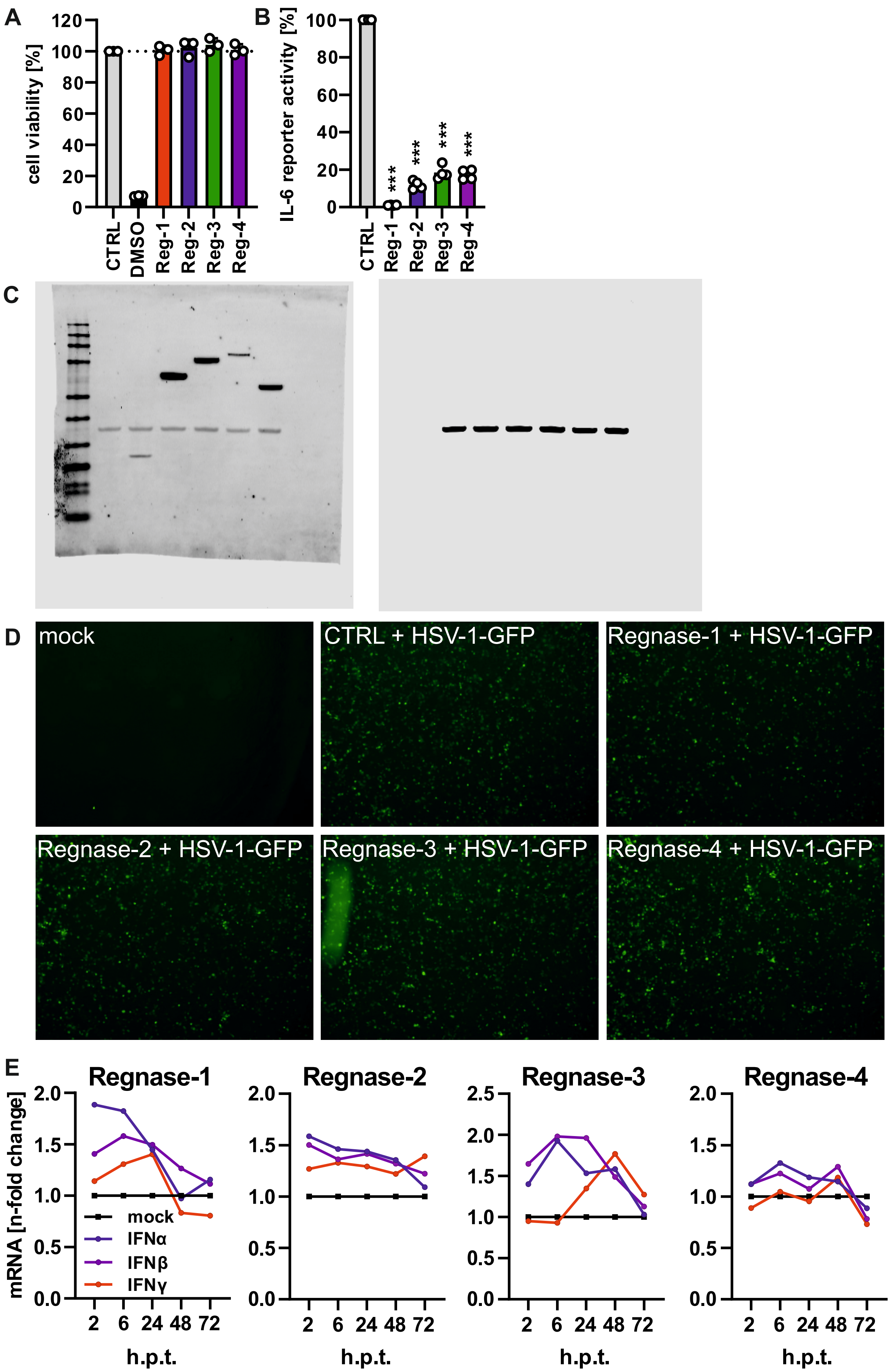

Grabe et al., 2026
Figure S2

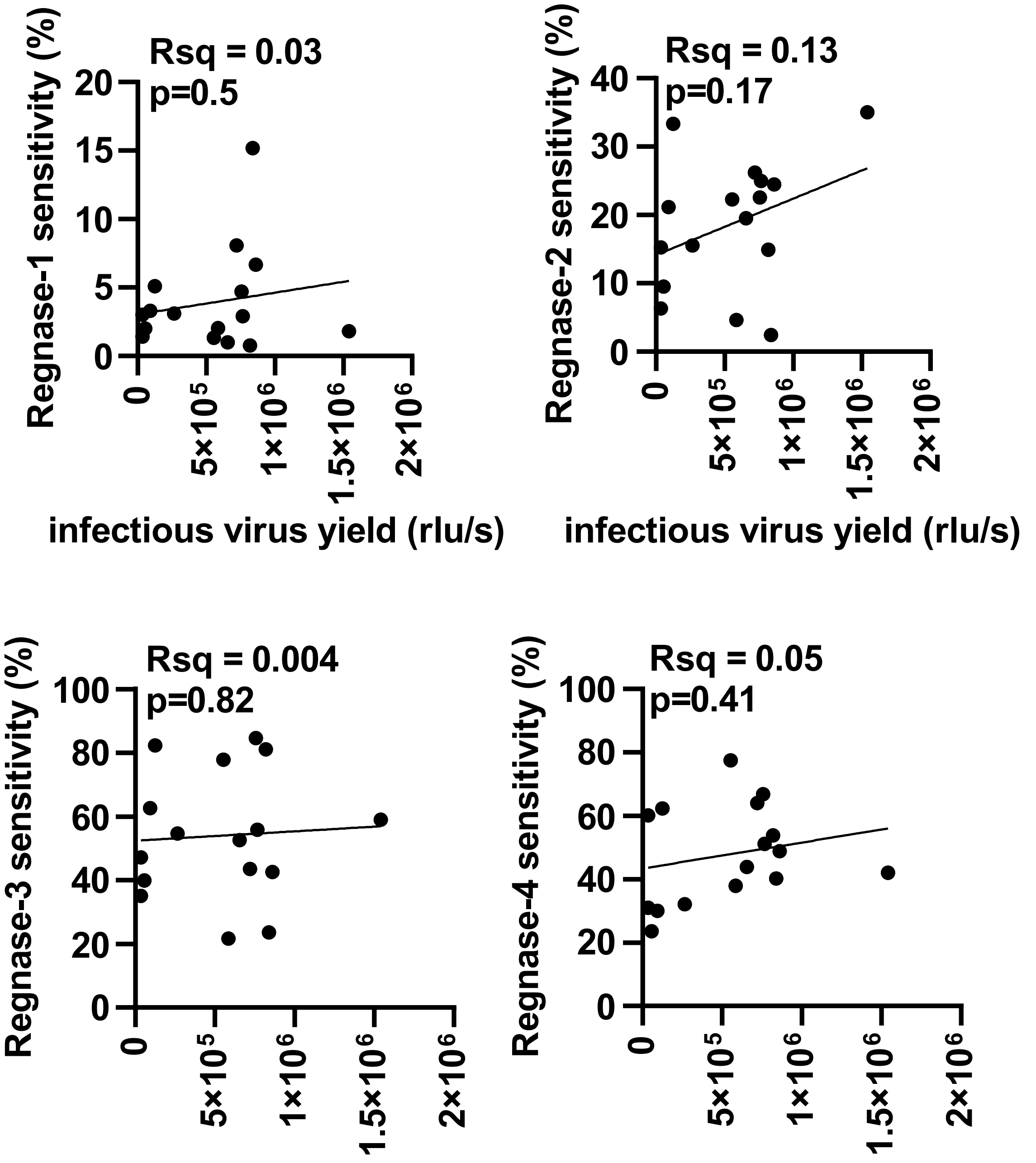

Grabe et al., 2026
Figure S3

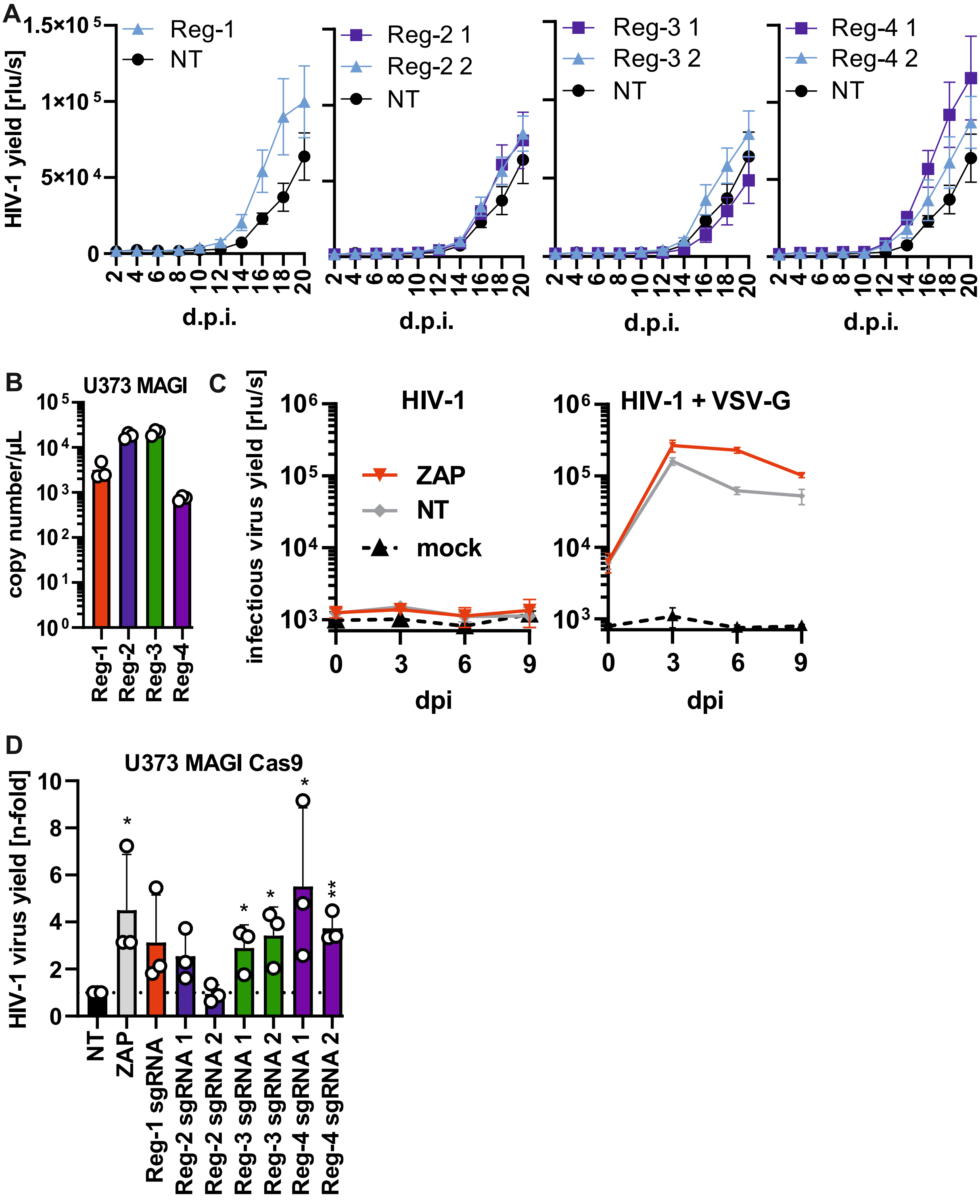

**Grabe et al., 2026**
**Figure S4**

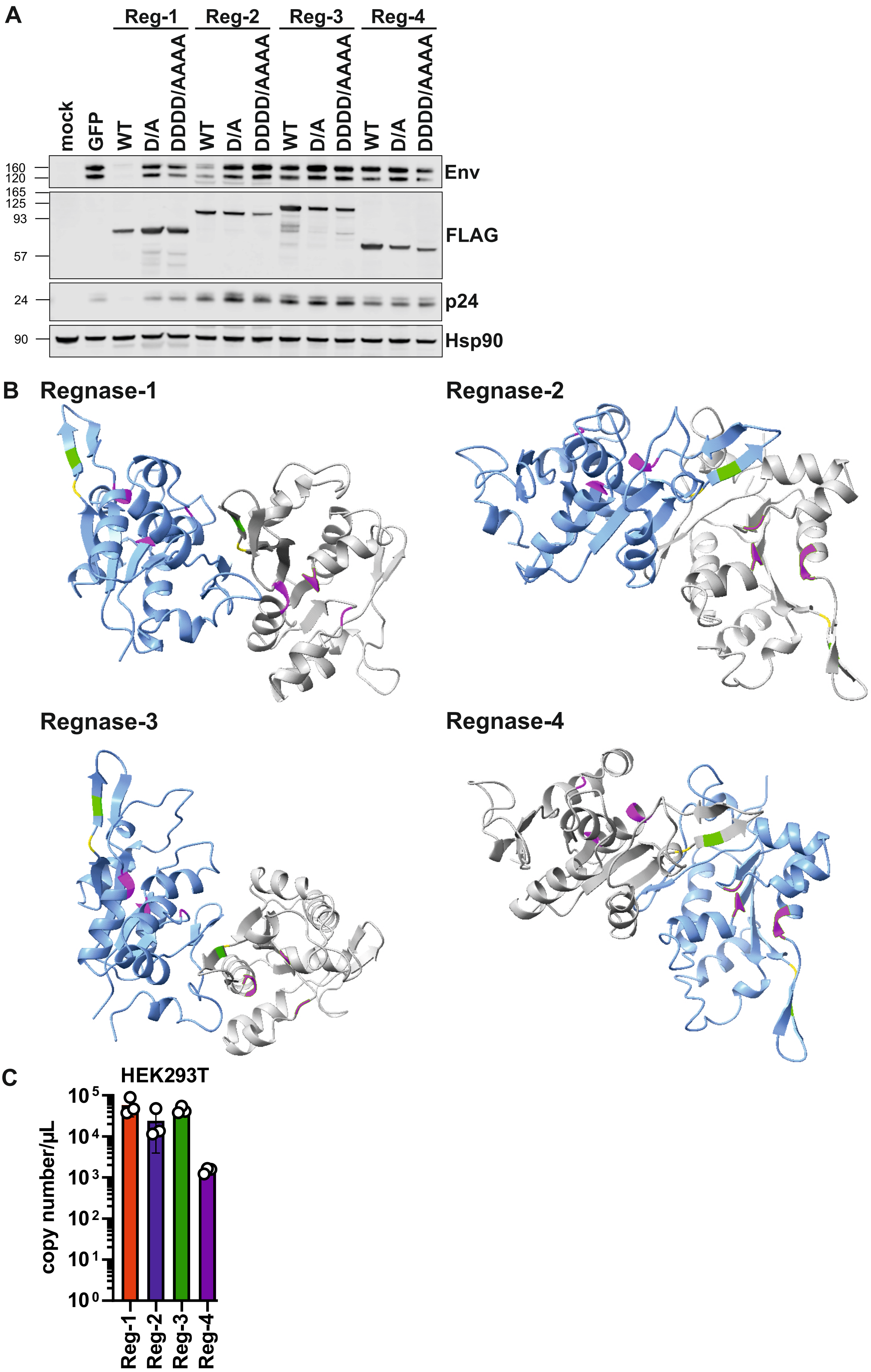

Grabe et al., 2026
Figure S5

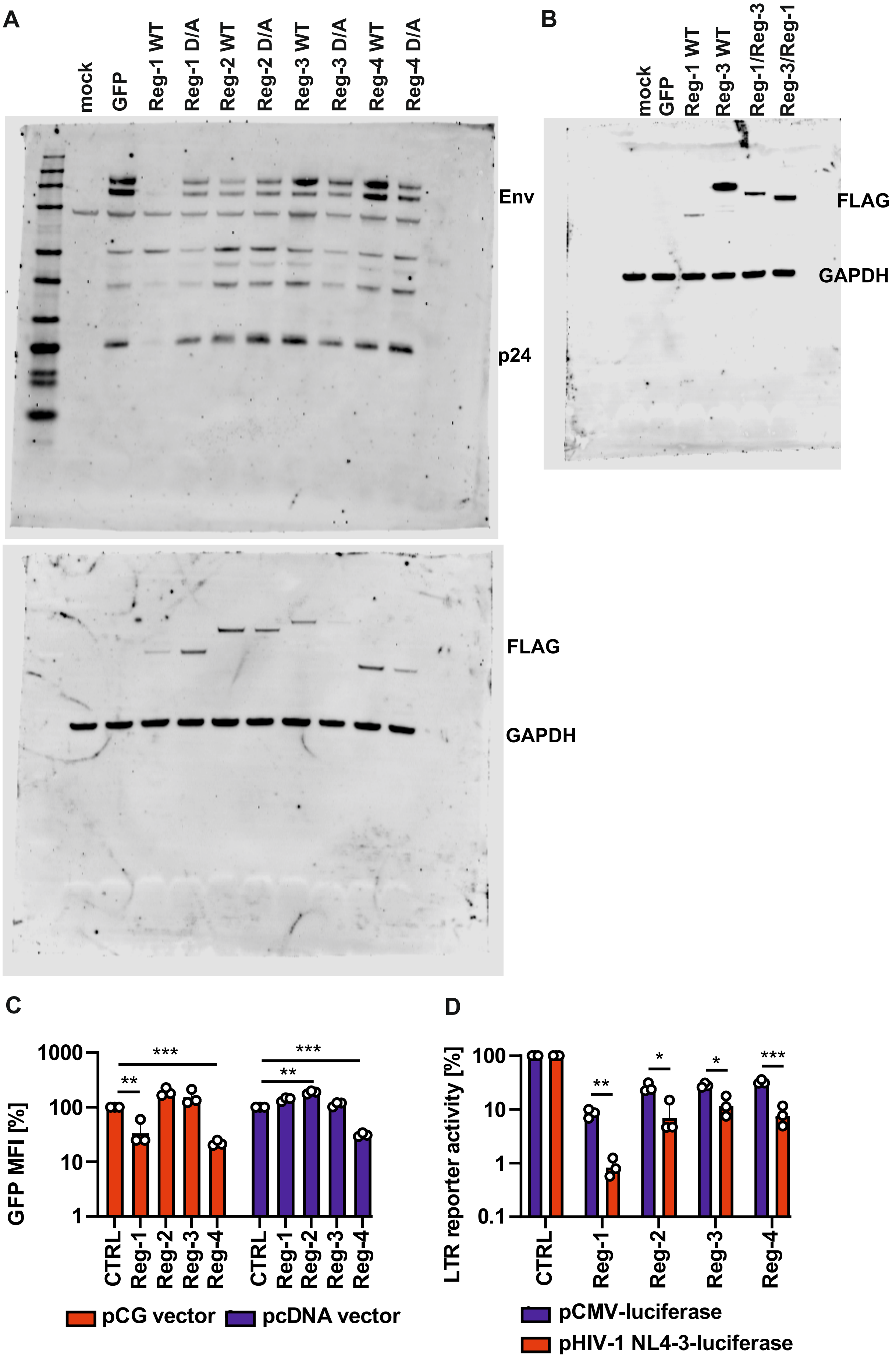
